# Dopamine Prediction Error Signaling in a Unique Nigrostriatal Circuit is Critical for Associative Fear Learning

**DOI:** 10.1101/2023.12.08.570564

**Authors:** Daphne Zafiri, Ximena I. Salinas-Hernández, Eloah S. De Biasi, Leonor Rebelo, Sevil Duvarci

## Abstract

Learning by experience that certain cues in the environment predict danger is crucial for survival. How dopamine (DA) circuits drive this form of associative learning is not fully understood. Here, we demonstrate that DA neurons projecting to a unique subregion of the dorsal striatum, the posterior tail of the striatum (TS), encode an aversive prediction error (PE) signal during associative fear learning. These DA neurons are necessary specifically during acquisition of fear learning, but not once the fear memory is formed, and are not required for forming cue-reward associations. Notably, temporally-precise excitation of DA terminals in TS is sufficient to enhance fear learning. Furthermore, neuronal activity in TS is crucial for acquisition of associative fear learning and learning-induced activity patterns in TS critically depend on DA input. Together, our results reveal that DA PE signaling in a non-canonical nigrostriatal circuit is crucial for driving associative fear learning.

## INTRODUCTION

Associative fear (threat; Ledoux, 2014) learning ─ the ability to associate stimuli with threats ─ enables animals to predict and avoid danger and hence is crucial for survival. However, learned fear that is excessive can also be maladaptive and have pathophysiological consequences. Much evidence indicates that anxiety disorders, such as post-traumatic stress disorder (PTSD), result from dysregulation of normal fear learning mechanisms (Craske et al., 2017; Duits et al., 2015; Shin et al., 2006). Therefore, elucidating the neural mechanisms underlying fear learning is critical for understanding the pathophysiology of anxiety disorders and thus has high clinical significance. In the laboratory, this kind of associative learning is commonly studied using Pavlovian fear conditioning (FC), in which an initially neutral stimulus (conditioned stimulus, CS) typically a tone is paired in time with an aversive unconditioned stimulus (US), such as a mild electrical foot shock. As the CS-US association is learned, the CS acquires the ability to elicit fear responses that are associated with the US (such as behavioral freezing) so that it can elicit conditioned fear responses when later presented alone. Traditionally, the amygdala, particularly its lateral nucleus (LA), has been recognized as the primary brain region for acquiring CS-US associations during FC (Duvarci and Pare, 2014; Johansen et al., 2011; LeDoux, 2000; Maren and Quirk, 2004; Sigurdsson et al., 2007; Tovote et al., 2015). However, recent studies suggest that plasticity in brain structures beyond the canonical amygdala circuitry is also involved in acquisition of fear memories. Of note, the posterior tail of the dorsal striatum (TS) has recently been implicated in fear conditioning (Kintscher et al., 2023; Chen et al., 2023).

TS is a unique subregion within the dorsal striatum that receives a combination of inputs from structures such as the auditory, visual and rhinal cortices as well as the amygdala, that sets it apart from other dorsal striatal subregions (Hunnicutt et al., 2016; Valjent and Gangarossa, 2021). Importantly, TS also receives dopaminergic innervation from a unique subpopulation of midbrain dopamine (DA) neurons that are predominantly located in the substantia nigra lateralis (SNL; Menegas et al., 2015). These DA neurons exhibit a distinct input-output organization compared to other midbrain DA neurons, project selectively to TS and do not overlap with the subpopulations that project to the rest of the striatum, cortex and the amygdala (Menegas et al., 2015). Notably, recent studies have shown that these DA neurons are particularly important for novelty-induced threat avoidance (Akiti et al., 2022; Menegas et al., 2018), and they have also been shown to be involved in fear learning (Chen et al., 2023). However, the precise role TS-projecting DA neurons play during associative fear learning is incompletely understood (Zafiri and Duvarci, 2022).

Associative learning is driven by prediction errors (PE) that signal the discrepancy between predicted and actual outcomes (Rescorla and Wagner, 1972); and new learning happens when outcomes do not match expectations. It is well-established that midbrain DA neurons, located in the ventral tegmental area (VTA) and the substantia nigra (SN), encode reward prediction errors (RPE) which act as teaching signals to drive reinforcement learning (Bayer and Glimcher, 2005; Eshel et al., 2015; 2016; Schultz et al., 1997; Steinberg et al., 2013). Notably, recent studies have further demonstrated that ventral midbrain DA neurons encode positive PE signals not only for rewards, but also for omission of aversive outcomes (Cai et al., 2020; Luo et al., 2018; Salinas-Hernández et al., 2018; 2023), to drive associative learning. However, although these DA neurons have been implicated in aversion (Verharen et al., 2020; Zafiri and Duvarci, 2022), their role in driving fear learning has remained elusive. Importantly, whether DA neurons projecting to brain structures outside the canonical amygdala circuitry contribute to PE signaling that is crucial for driving associative fear learning is largely unknown.

In this study, we investigate the precise role DA projections to TS play during associative fear learning. By performing measurements of DA terminal activity as well as DA release in TS, we found that DA projections to TS encode an aversive PE signal during associative fear learning.

Selective lesioning of TS-projecting DA neurons demonstrated that these neurons are required specifically during acquisition of fear learning, but not once the CS-US association was learned. Notably, activity of these DA neurons was crucial for associating sensory cues with the aversive US, but not the context with aversive US or the sensory cues with reward. Conversely, temporally-precise excitation of DA terminals in TS during FC was sufficient to enhance fear learning. To gain further insights into the functional role of TS, we performed Ca^2+^ recordings of TS activity and found a PE-like activity pattern, as well as potentiation of CS responses, during associative fear learning. Bidirectional manipulations of TS activity further showed that the neuronal activity in TS is required for and sufficient to enhance fear learning. Finally, we demonstrated that DA input was crucial for the fear learning-induced activity patterns in TS during FC. Taken together, our results reveal a key role for DA PE signaling in a unique nigrostriatal circuit for driving associative fear learning.

## RESULTS

### DA Neurons Projecting to TS Encode an Aversive PE Signal during Associative Fear Learning

In order to investigate the activity of DA neurons projecting to TS in a projection-specific manner, we used fiber photometry to measure activity-dependent Ca^2+^ signals at the terminals of DA neurons in TS. We injected a Cre-dependent adeno-associated virus (AAV) expressing the genetically encoded Ca^2+^ indicator GCaMP in SN of transgenic mice expressing Cre recombinase under the control of the dopamine transporter (DAT) promoter (DAT-Cre mice; Figure 1A). In these mice, the expression of Cre is highly selective for DA neurons (Lammel et al., 2015). In line with this, we observed a high degree of overlap between Cre-dependent GCaMP6m expression and immunohistochemical staining against tyrosine hydroxylase (TH; Figure 1C) in the SN. An optical fiber implanted in TS (Figure 1A-B, Supplementary Figure 1) enabled recording of Ca^2+^ transients in the axon terminals of DA neurons (Figure 1D).

**Figure 1.**
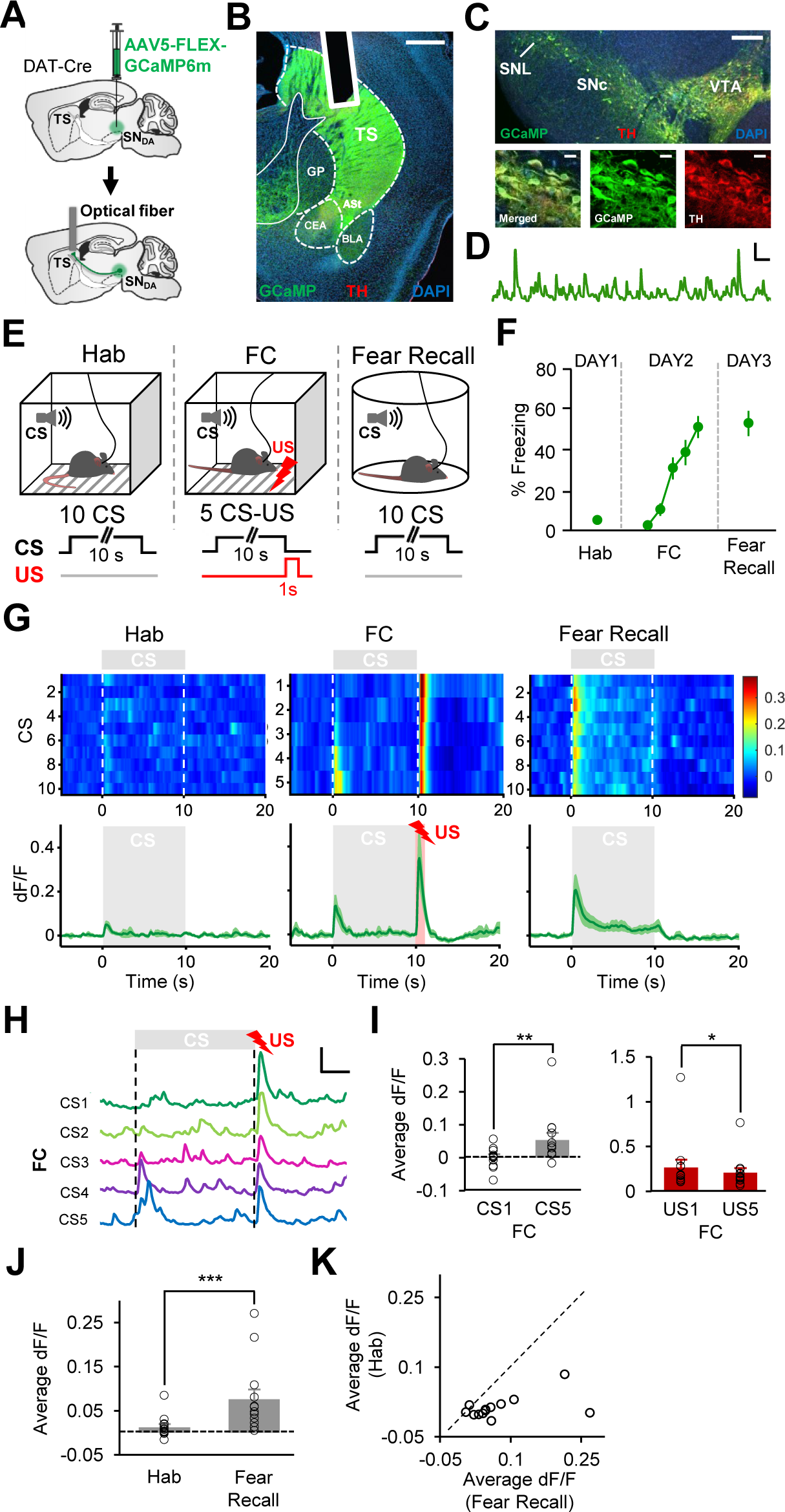
DA terminals in TS encode an aversive PE signal during associative fear learning. A) Schematic of the surgical procedure showing virus injection in SN (top) and optical fiber implantation in TS (bottom). **B)** Example histological image showing expression of GCaMP (green) along with immunostaining for tyrosine hydroxylase (TH, red) at DA terminals and DAPI (blue) staining in TS. White vertical track indicates the optical fiber placement in TS. ASt, amygdalostriatal transition area; BLA, basolateral amygdala; CEA, central nucleus of the amygdala; GP, globus pallidus. Scale bar: 0.5 mm. **C)** Top: example histological image showing Cre-dependent expression of GCaMP (green) along with immunostaining for TH (red) and DAPI (blue) staining. SNc, substantia nigra pars compacta; SNL, substantia nigra lateralis; VTA, ventral tegmental area. Scale bar: 0.25 mm. Bottom: confocal images showing expression of GCaMP and TH staining and the merged image showing co-expression. Scale bar: 20 μm. **D)** Example of change in fluorescence (dF/F) over time in the axon terminals of DA neurons in TS. Scale bar: 5s, 0.5 dF/F. **E)** Top: schematic of the behavioral protocol. Hab: tone habituation, FC: fear conditioning. Bottom: schematic of CS and US presentations during Hab, FC and Fear Recall sessions. **F)** Behavioral freezing to the CS (n = 13 mice) during Hab. (average of 10 CSs), FC and Fear Recall sessions (average of 10 CSs). Mice froze significantly higher to the CS at the end of FC (t-test, t(12) = 9.22, p < 0.0001) and during Fear Recall (t-test, t(12) = 8.78, p < 0.0001) compared to Hab. **G)** Top: Average activity of DA terminals around each CS during Hab, FC and Fear Recall across all mice (n = 13). The heat map shows response amplitudes (dF/F) around each CS presentation. Bottom: Average change in fluorescence around the time of CS presentation (gray area) during Hab, FC and Fear Recall. The red area during FC represents the US presentation. **H)** Change in fluorescence around each CS and US presentation during FC in an example animal. Scale bar: 2.5s, 0.5 dF/F. **I)** Left: comparison of average change in fluorescence in the 5s after CS onset during CS1 and CS5 of FC. Note the significant increase in the Ca^2+^ signal from CS1 to CS5 (**p = 0.0017, signed-rank test). Right: comparison of average change in fluorescence during the US (1s) for US1 and US5 of FC. Note the significant decrease in the Ca^2+^ signal from US1 to US5 (*p = 0.013, signed-rank test). **J)** Average change in fluorescence in the 5s after CS onset during Hab and Fear Recall. Note the significant increase in the Ca^2+^ signal from Hab to Fear Recall (***p = 0.0007, signed-rank test). **K)** Scatter plot showing CS responses of each recording site during Hab and Fear Recall. Data points below the unity line represent larger CS responses during Fear Recall. Shaded regions and error bars represent mean ± s.e.m across animals.

To examine DA terminal activity during associative fear learning, mice (n = 13) were trained in a FC paradigm (Figure 1E) where a tone (CS) was paired with an aversive foot shock (US) on day 2, following a tone habituation session (Hab) on day 1. On day 3, mice received a fear recall session consisting of CS presentations in the absence of the aversive US. During the course of FC, freezing to the CS gradually increased (Figure 1F), indicating that the mice learned the association between the CS and the US. Mice showed significantly higher freezing levels to the CS at the end of FC session, as well as during fear recall, compared to Hab (Figure 1F). Notably, the activity of DA terminals in TS appeared to resemble a PE signal during associative fear learning. In the beginning of FC, DA terminals in TS showed strong excitation to the aversive US which decreased during the course of conditioning (Figure 1G-I), as the CS-US association was learned and the CS came to predict the occurrence of the US. Indeed, there was a significant decrease in US responses from the first to the last US (Figure 1I). Conversely, while CS responses were absent at the beginning of FC, they gradually increased through the course of conditioning (Figure 1G-I), mirroring the increase in freezing to the CS (Figure 1F). In line with the behavioral results, responses to the CS were significantly larger during fear recall compared to Hab session (Figure 1J), indicating that the CS responses were potentiated as the animals learned the CS-US association. In almost all animals, responses to the CS were larger during fear recall compared to Hab session (Figure 1K). Together, these results demonstrated that DA terminals in TS exhibited a PE-like activity pattern during associative fear learning and suggested that TS-projecting DA neurons encode an aversive PE signal.

Because presentation of the aversive US itself could result in a nonspecific enhancement of CS responses, we next asked whether potentiation of CS responses during FC was indeed a result of associative learning and depended on the temporally contingent presentations of the CS and the US. To address this question, mice underwent an unpaired training paradigm where they received same number of CS and US presentations as in FC, but the CS and the US were explicitly unpaired in time during the training session on day 2 (Figure 2A). In mice undergoing the unpaired training, we did not observe an increase in freezing to the CS between Hab, training and testing sessions (Figure 2B). Mirroring these behavior results, we also did not find a significant difference in the CS response between the Hab and the testing session (n = 7; p = 0.31, signed-rank test Figure 2C-E). These results indicate that potentiation of CS responses as well as the increased behavioral fear responses to the CS during FC required the association of the CS with the aversive US.

**Figure 2.**
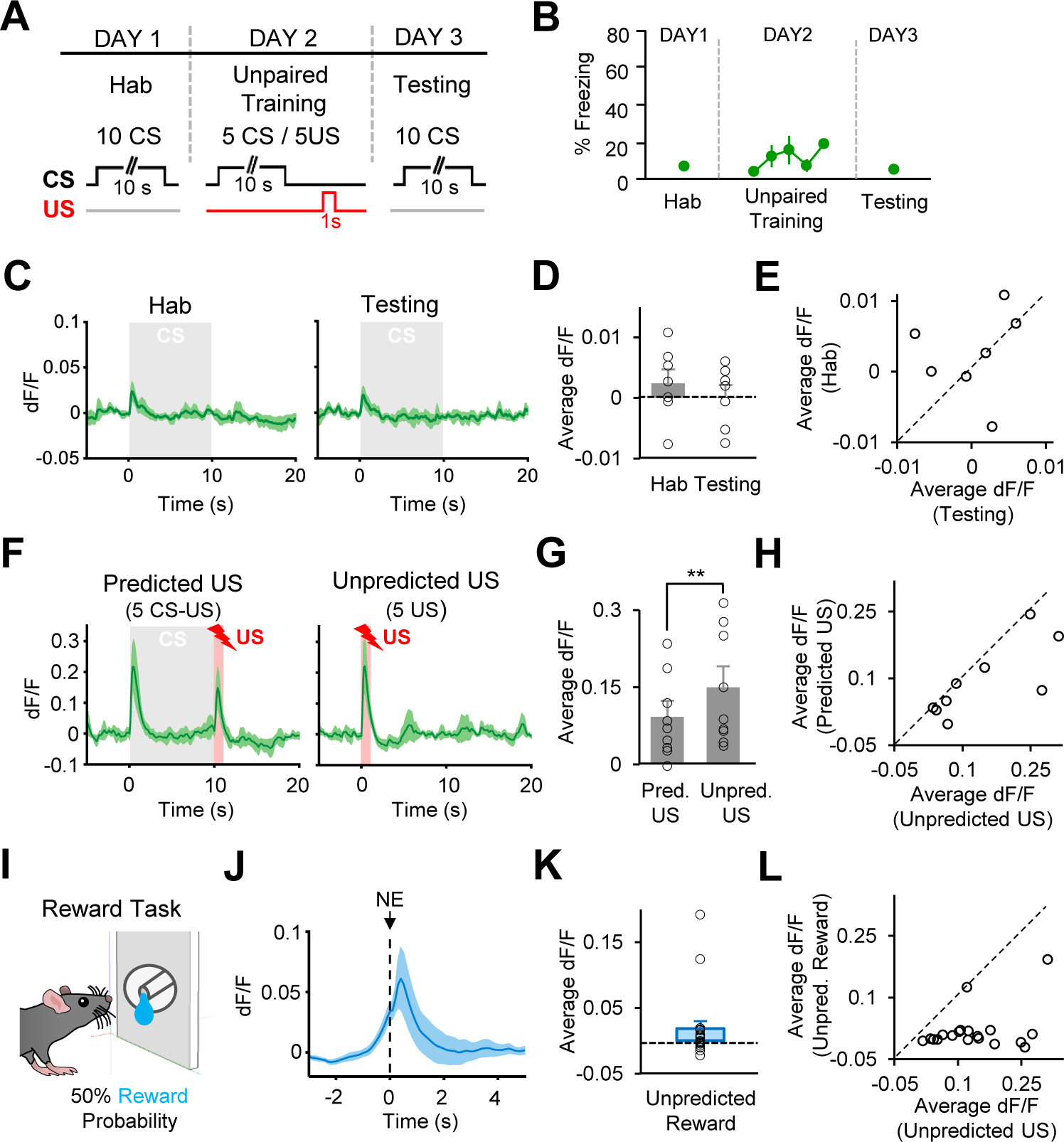
Activity of DA terminals in TS in response to CSs, aversive USs and rewards that are unpredicted. A) Top: schematic of the behavioral protocol for unpaired training. Hab: tone habituation. Bottom: schematic of CS and US presentations during Hab, Unpaired Training and Testing sessions. **B)** Behavioral freezing to the CS (n = 6 mice) during Hab. (average of 10 CSs), Unpaired Training and Testing sessions (average of 10 CSs). **C)** Average change in fluorescence from recording sites in TS (n = 7) around the time of CS (gray area) during Hab, and Testing. **D)** Comparison of average change in fluorescence in the 5s after CS onset during Hab. and Testing (p = 0.31, signed-rank test). **E)** Scatter plot showing CS responses of each recording site during Hab and Testing. Data points below the unity line represent larger CS responses during Testing. **F)** Average change in fluorescence from recording sites in TS (n = 9) around the time of CS (gray area) and US (red area) during Predicted and Unpredicted US presentations. **G)** Comparison of average change in fluorescence during 1s US presentation for Predicted and Unpredicted USs (**p = 0.0078, signed-rank test). **H)** Scatter plot showing US responses of each recording site during Predicted and Unpredicted US presentations. Data points below the unity line represent larger responses for Unpredicted US. **I)** Schematic of the reward task. Animals received reward 50% of the time after entering the noseport. **J)** Average change in fluorescence during rewarded noseport entries (NE) from all recording sites (n = 20). **K)** Average change in fluorescence in the 3s after noseport entry during rewarded NE. **L)** Scatter plot showing responses from each recording site (n = 20) during Unpredicted US and reward presentations. Data points below the unity line represent larger responses for Unpredicted US. Shaded regions and error bars represent mean ± s.e.m across animals.

If DA terminals in TS encode an aversive PE signal, we expect that responses to unpredicted USs should be larger in magnitude compared to responses to predicted USs. To test this, we examined DA terminal activity while previously well-trained mice received presentations of predicted (CS-US pairings) and unpredicted (US only) foot shocks (Figure 2F). We indeed found stronger responses to unpredicted USs compared to CS-predicted ones (n = 9; p = 0.0078, signed-rank test; Figure 2G, H), consistent with the decrease in US responses that we observed during the course of FC. Together, these results indicate that TS-projecting DA neurons signal a PE for aversive outcomes.

Our results so far demonstrate that TS-projecting DA neurons are strongly activated by aversive USs. An important question is whether these DA neurons encode the motivational salience of stimuli. If they signal salience, we expect that they would also be activated strongly by rewards. Since DA neurons are known to respond to rewards particularly when they are unpredicted (Bayer and Glimcher, 2005; Eshel et al., 2015; 2016; Roesch et al., 2007; Schultz et al., 1997), we used a reward task in which mice nose-poked to receive rewards with 50% probability (Figure 2I), making reward delivery unpredicted. Interestingly, we found that DA terminals in TS exhibited weak responses to rewards (n = 20; Figure 2J, K), consistent with previous reports (Menegas et al., 2017; 2018). In all animals tested, responses to unpredicted rewards were much smaller compared to unpredicted footshock USs (Figure 2L). Together, our results indicated that while DA terminals in TS were strongly activated particularly by painful and aversive outcomes, they responded only weakly to rewards, suggesting that these neurons likely did not encode the motivational salience of stimuli.

### DA Release in TS Signals an Aversive PE during Associative Fear Learning

DA neurons have been shown to co-release glutamate and GABA in the striatum (Adrover et al., 2014; Tritsch et al., 2012). It is therefore possible that the DA terminal activity might not reflect DA release during associative fear learning. To address this, we next examined the dynamics of DA release in TS during associative fear learning by performing optical recordings of the genetically encoded DA biosensor dLight (Patriarchi et al., 2018) using fiber photometry. To this end, an AAV expressing dLight (AAV5-CAG-dLight1.1) was injected and optical fibers were implanted in the TS (Figure 3A-B, Supplementary Figure 2). Mice (n = 8) underwent the same fear conditioning protocol as in the GCaMP recording experiment (Figure 3C), and showed increased freezing to the CS following FC (Figure 3D), indicating successful fear learning.

**Figure 3.**
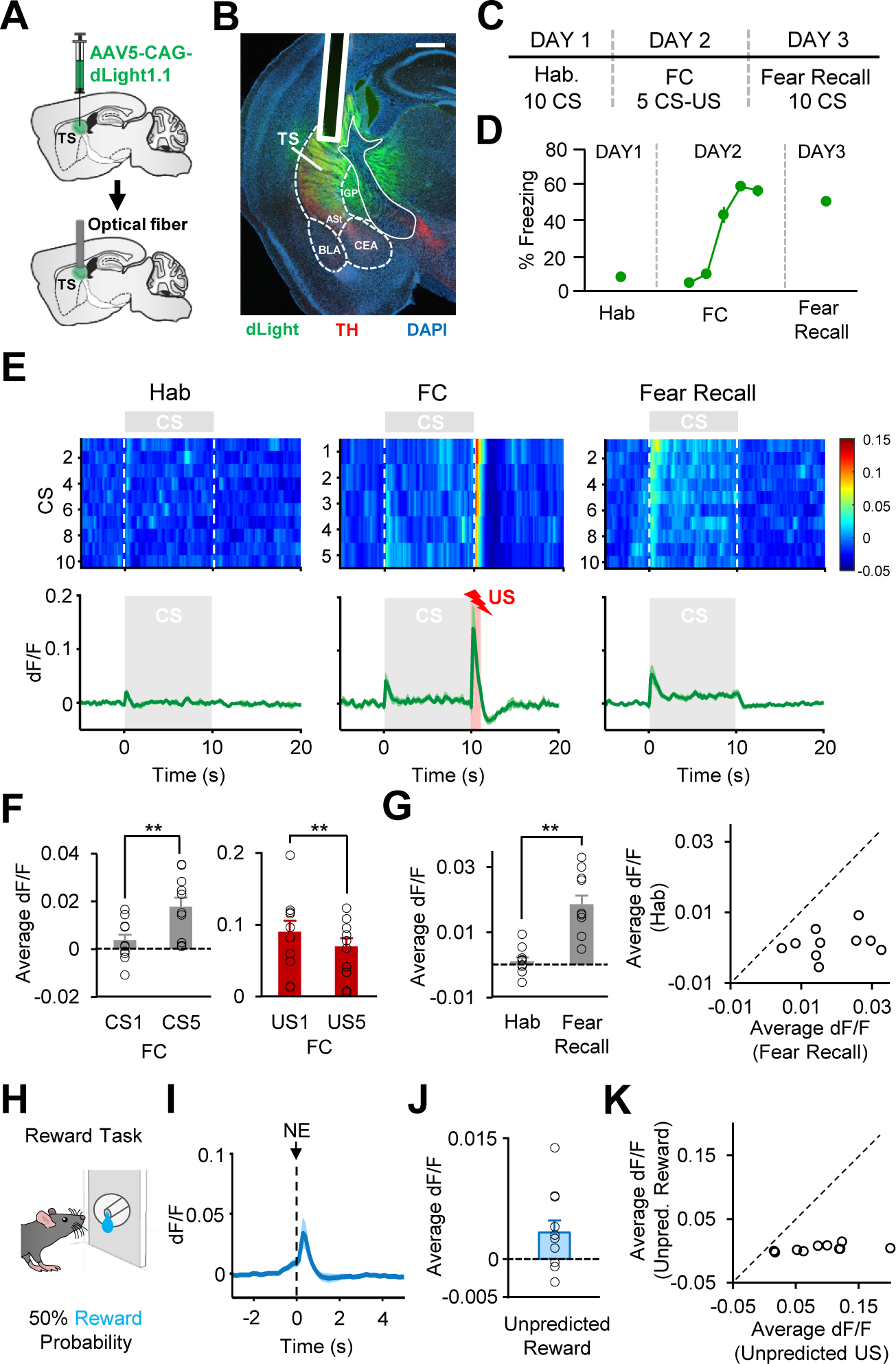
DA release dynamics in TS during associative fear learning and reward task. A) Schematic of the surgical procedure showing virus injection (top) and optical fiber implantation (bottom) in TS. **B)** Example histological image showing expression of dLight (green) along with immunostaining for tyrosine hydroxylase (TH, red) and DAPI (blue) staining in TS. White vertical track indicates the optical fiber placement in TS. ASt, amygdalostriatal transition area; BLA, basolateral amygdala; CEA, central nucleus of the amygdala; GP, globus pallidus. Scale bar: 0.5 mm. **C)** Schematic of the behavioral protocol. Hab: tone habituation, FC: fear conditioning. **D)** Behavioral freezing to the CS (n = 8 mice) during Hab, FC and Fear Recall. **E)** Top: Average dLight activity around each CS during Hab, FC and Fear Recall across all recording sites (n = 10). The heat map shows response amplitudes (dF/F) around each CS presentation. Bottom: Average change in fluorescence around the time of CS presentation (gray area) during Hab, FC and Fear Recall. The red area during FC represents the US presentation. **F)** Left: comparison of average change in fluorescence in the 5s after CS onset during CS1 and CS5 of FC. Note the significant increase in the dLight signal from CS1 to CS5 (**p = 0.008, signed-rank test). Right: comparison of average change in fluorescence during the US (1s) for US1 and US5 of FC. Note the significant decrease in the dLight signal from US1 to US5 (**p = 0.009, signed-rank test). **G)** Left: Average change in fluorescence in the 5s after CS onset during Hab and Fear Recall. Note the significant increase in the dLight signal from Hab to Fear Recall (**p = 0.002, signed-rank test). Right: scatter plot showing CS responses of each recording site (n = 10) during Hab and Fear Recall. Data points below the unity line represent larger CS responses during Fear Recall. Shaded regions and error bars represent mean ± s.e.m across animals. **H)** Schematic of reward task. Animals received reward 50% of the time after entering the noseport. **I)** Average change in fluorescence during rewarded noseport entries (NE). **J)** Average change in fluorescence in the 3s after noseport entry during rewarded NE. **K)** Scatter plot showing responses from each recording site (n = 10) during unpredicted US (first US of FC) and reward. Data points below the unity line represent larger responses for Unpredicted US. Shaded regions and error bars represent mean ± s.e.m across animals.

In line with Ca^2+^ recordings in DA terminals, we found strong dLight responses to the US at the beginning of FC which significantly decreased through the course of conditioning (Figure 3E, F), as occurrence of the US became predictable. Conversely, we observed a significant increase in the dLight activity in response to the CS when the first and last CSs of FC were compared (Figure 3E, F), indicating potentiation of CS responses as the CS-US association was learned. Consistent with increased behavioral freezing, CS responses exhibited a significant increase from Hab to fear recall (Figure 3E, G), and in all recording sites (n = 10) CS responses during fear recall were larger compared to Hab (Figure 3G). Furthermore, we again observed small responses to unpredicted rewards (Figure 3H-J), and in all dLight recording sites unpredicted footshock US responses (responses to the first US of FC) were larger compared to unpredicted reward responses (Figure 3K). Taken together, these results indicate that DA release in TS underlies PE signaling during associative fear learning.

### TS-projecting DA Neurons are Required Selectively for Acquisition of Cued Associative Fear Learning

PE signals are thought to drive associative learning (Rescorla and Wagner, 1972). If DA neurons projecting to TS encode an aversive PE signal and this signal is critical for driving associative fear learning, we expect that lesioning these DA neurons should impair particularly the acquisition of fear conditioning. To address this, we performed projection-specific ablation of TS-projecting DA neurons using a DA neuron selective neurotoxin, 6-hydroxydopamine (6- OHDA; Figure 4A). Importantly, the 6-OHDA injections in TS caused reduction of DA axons specifically in TS, and not in the neighboring structures such as the amygdala and the amygdalostriatal transition area (ASt; Figure 4B). Following 6-OHDA lesions, mice were trained using an auditory FC protocol (Figure 4C) consisting of 5 CS-US pairings. Twenty-four hours after conditioning, mice underwent a fear recall test consisting of 5 CS presentations. We found that the lesioned mice (n = 10) froze significantly less to the CS throughout the conditioning session compared to saline-injected control mice (n = 12; Figure 4D), suggesting impaired fear learning. Furthermore, impaired fear acquisition resulted in a weaker fear memory when tested the next day (Figure 4D). 6-OHDA lesioned mice froze significantly less at the end of FC and the beginning of fear recall sessions compared to control mice (Figure 4E). These results demonstrated that TS-projecting DA neurons are required for acquisition of the CS-US association.

**Figure 4.**
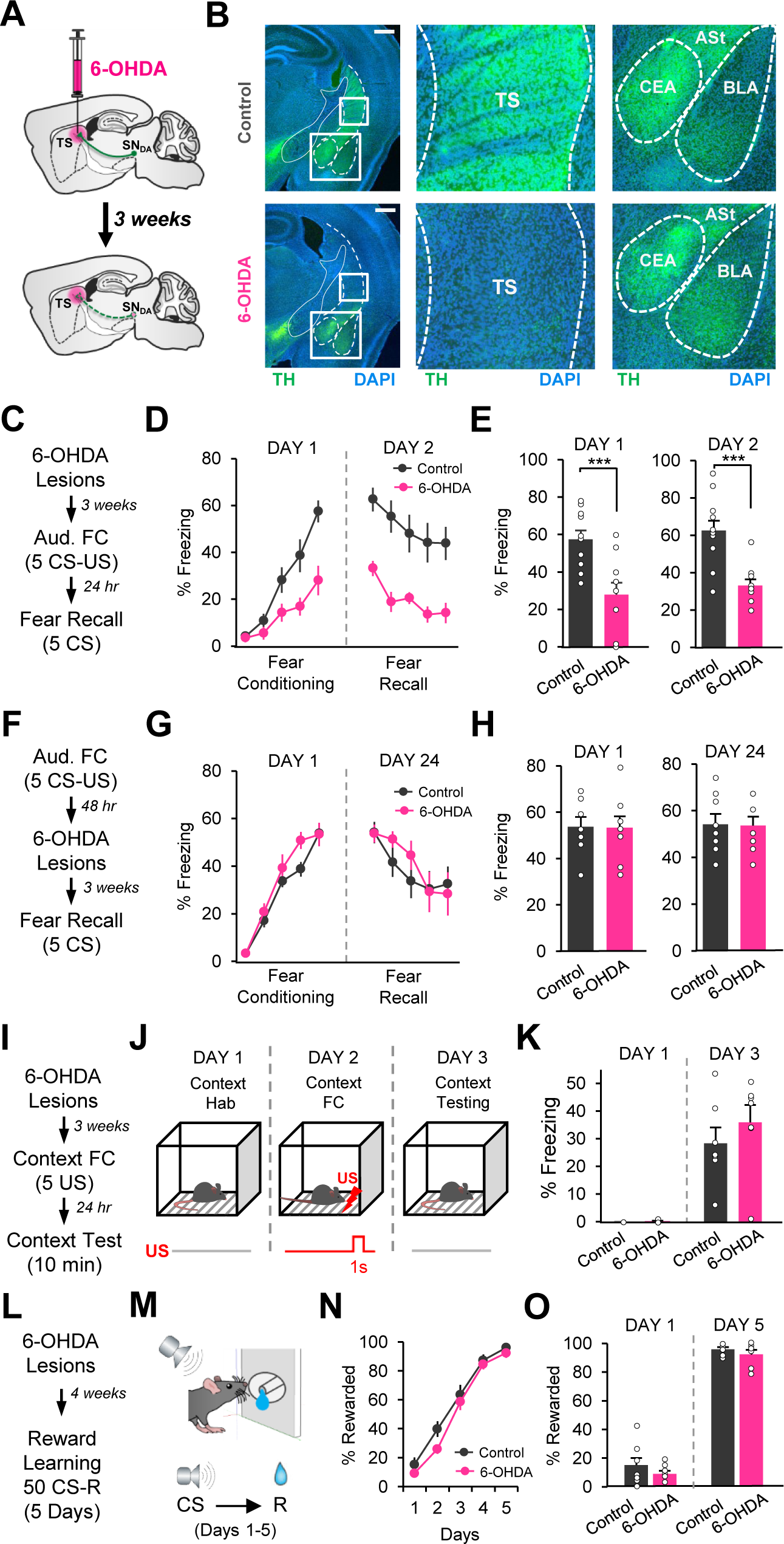
TS-projecting DA neurons are required selectively for acquisition of cued associative fear learning. A) Schematic of the surgical procedure showing 6- OHDA injections in TS (top) and ablation of TS-projecting DA neurons in the SN (SN_DA_) three weeks later (bottom). **B)** Example coronal sections showing immunostaining for tyrosine hydroxylase (TH, green) and DAPI (blue) staining in control saline (top, left) and 6-OHDA (bottom, left) injected animals. Scale bar: 0.5 mm. Close up images showing TH-staining in TS (middle) and the amygdala (right; BLA, basolateral amygdala; CEA, central amygdala) in saline (top) and 6-OHDA (bottom) injected animals. Note the reduction in TH immunostaining in TS, but not in the neighboring structures such as the amygdala and the amygdalostriatal transition area (Ast), in 6-OHDA injected mice. **C)** Schematic of the behavioral protocol for 6-OHDA ablation before fear acquisition. Aud. FC: auditory fear conditioning. **D)** Behavioral freezing to the CS during FC and fear recall test for mice that received saline (n = 12) and 6-OHDA (n = 10) injections before fear conditioning. 6-OHDA group showed impaired fear acquisition and fear recall (two-way repeated measures ANOVA, fear conditioning: main effect of group: F_1,80_ = 7.84, p = 0.011; group × trial interaction: F_4,80_ = 7.81, p < 0.0001; fear recall: main effect of group: F_1,80_ = 18.60, p = 0.0003) **E)** Left: freezing to the last CS during fear conditioning. 6-OHDA group froze significantly less compared to saline-injected control group (t-test, t(20) = 3.85, p = 0.001). Right: freezing to the CS at the beginning of fear recall (first CS). 6-OHDA group froze significantly less compared to control group (t-test, t(20) = 4.68, p = 0.0001). **F)** Schematic of the behavioral protocol for 6-OHDA ablation after fear acquisition. **G)** Freezing to the CS during FC and fear recall test for mice that received saline (n = 8) and 6-OHDA (n = 8) injections after fear conditioning (two-way repeated measures ANOVA, fear conditioning: no main effect of group: F_1,56_ = 1.37, p = 0.26 and no group × trial interaction: F_4,56_ = 1.39, p = 0.24; fear recall: no main effect of group: F_1,56_ = 0.12, p = 0.72 and no group × trial interaction: F_4,56_ = 1.52, p = 0.20). **H)** Left: freezing to the last CS during FC (t-test, t(14) = 0.06, p = 0.95). Right: freezing to the CS at the beginning of fear recall (first CS; t-test, t(14) = 0.07, p = 0.94). **I)** Schematic of the behavioral protocol for 6-OHDA ablation before contextual fear acquisition. **J)** Schematic of contextual fear conditioning task. Hab: habituation. **K)** Freezing to the context during Context Hab. and Context Test in control (n = 7) and 6-OHDA (n = 7) groups. The two groups showed comparable freezing levels during Context Hab (p = 0.14, t-test) and Context Test (p = 0.38, t-test), indicating 6-OHDA ablation of TS- projecting DA neurons does not have an effect on contextual fear learning. **L)** Schematic of the behavioral protocol for 6-OHDA ablation before reward learning. **M)** Schematic of the reward learning task. **N)** Percent of rewarded CSs over the course of the reward task. Control (n = 8) and 6-OHDA (n = 7) groups showed comparable performance (two-way repeated measures ANOVA, no main effect of group: F_1,52_ = 1.7, p = 0.21 and no group × trial interaction: F_4,52_ = 0.93, p = 0.45). **O)** Percent of rewarded CSs during the first and last days of the reward task. The two groups exhibited comparable performance. Error bars represent mean ± s.e.m. across animals.

An aversive PE signal is expected specifically to be critical for initiating and driving new associative fear learning, but not the retrieval of fear memories. We therefore hypothesized that TS-projecting DA neurons might be crucial selectively during fear acquisition on the conditioning day, but not later once the CS-US association was learned. To test this, we performed 6-OHDA ablation of TS-projecting DA neurons after fear memory was formed (Figure 4F). In support of our hypothesis, we found that the lesioned mice exhibited comparable levels of freezing to the CS during the fear recall test performed 3 weeks after 6-OHDA lesions (Figure 4G, H), suggesting that the activity of these DA neurons was not necessary for retrieval and expression of fear memories. These results indicated that TS-projecting DA neurons were indeed crucial selectively for acquisition of fear conditioning, but were no longer required once the CS-US association was learned.

Given that TS is a sensory striatal region receiving mainly auditory and visual inputs (Hunnicutt et al., 2016), we hypothesized that DA projections to TS might be important for associating specifically the sensory cues with aversive outcomes. To address this, we next investigated whether TS-projecting DA neurons were necessary for learning the association between the context and the aversive US. To this end, we performed a contextual FC paradigm consisting of 5 US presentations in context A. The animals were tested for contextual fear memory the next day (Figure 4J). Mice received 6-OHDA or saline injections in TS as described above and 3 weeks later underwent contextual FC (Figure 4I). Lesioned and control mice showed comparable levels of freezing to the context during contextual testing (Figure 4K), indicating that DA projections to TS were not required for learning the association between the context and the US.

Since we observed only small responses to rewards, we hypothesized that TS-projecting DA neurons might not be required for learning the association between a cue and reward. To test this, we performed a Pavlovian reward learning task (Figure 4L, M) in which a tone CS was paired with reward. The animals underwent the reward task following 6-OHDA lesioning of TS-projecting DA neurons (Figure 4L). Notably, both lesioned and control groups showed similar learning rates during the reward task (Figure 4N, O), suggesting that TS-projecting DA neurons are not required for associating cues with rewards and hence are not critical for Pavlovian reward learning.

Taken together, these results reveal a highly selective role for TS-projecting DA neurons in associative learning. We demonstrate that they are crucial selectively during the acquisition of fear learning, but not once the CS-US association is learned. Importantly, we found that these DA neurons are specifically required for learning auditory CS−US, but not context−US or auditory CS−reward, associations.

### Temporally-Precise Activation of DA Terminals in TS is Sufficient to Enhance Associative Fear Learning

If DA activity in TS at the time of the US acts as a teaching signal and causes learning about the CS, then boosting this signal should enhance associative fear learning. To test this, we optogenetically excited DA terminals in TS precisely at the time of the US during FC. DAT-Cre mice were bilaterally injected with a Cre-dependent AAV expressing either channelrhodopsin-2 (ChR2) fused with EYFP (ChR2-EYFP) or EYFP only (EYFP control) in the SN, and implanted bilaterally with optical fibers above TS (Figure 5A-C, Supplementary Figure 3). We again observed a high level of overlap between Cre-dependent ChR2-EYFP expression and immunohistochemical staining against TH (Figure 5C) suggesting DA neuron-specific expression of ChR2.

**Figure 5.**
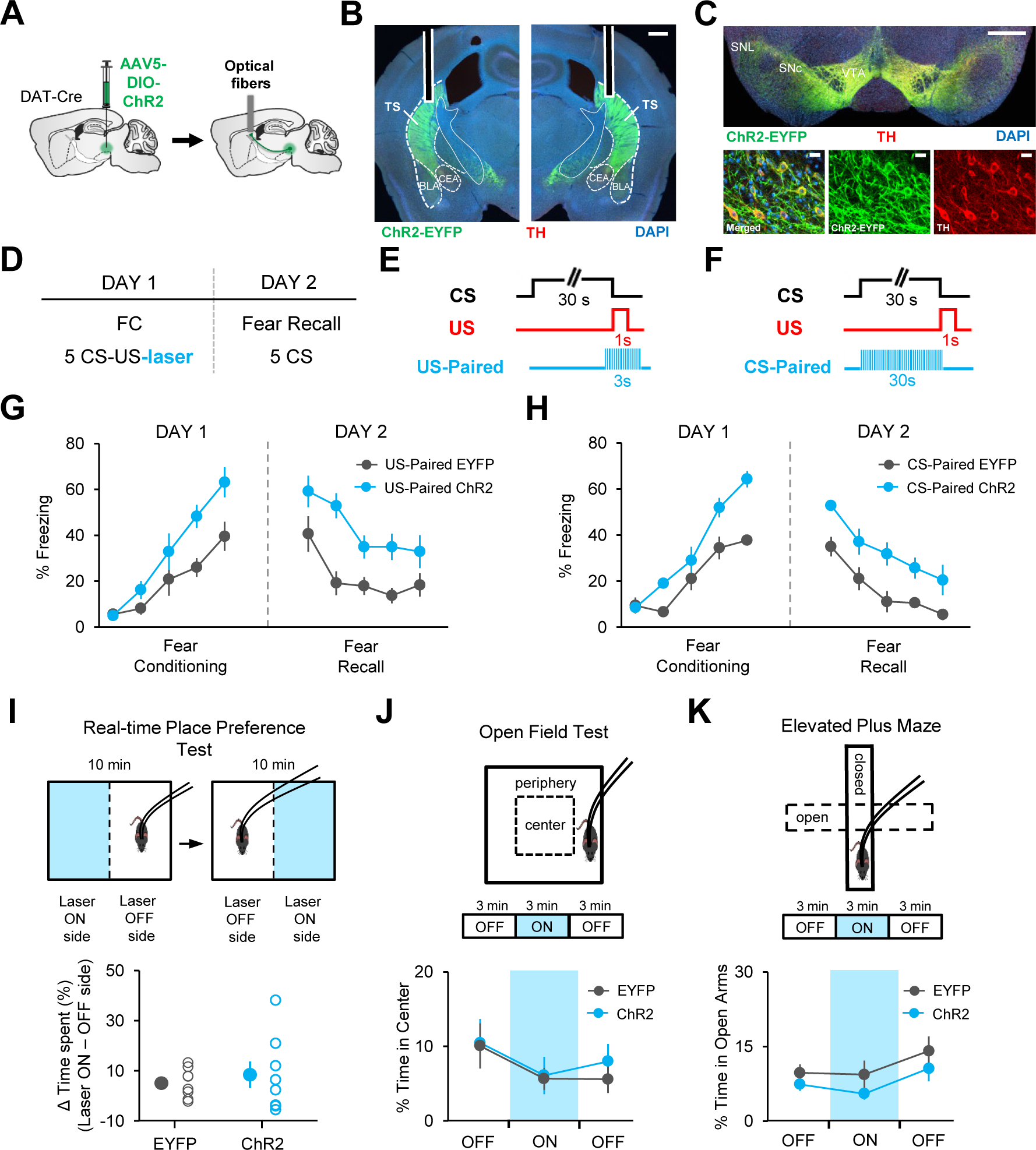
Temporally-precise activation of DA terminals in TS is sufficent to enhance associative fear learning. A) Schematic of the surgical procedure showing virus injection in the SN and optical fiber implantation in TS. **B)** Example histological image showing optical fiber placements and Cre-dependent expression of ChR2-EYFP at DA terminals (green) along with immunostaining for tyrosine hydroxylase (TH, red) and DAPI (blue) staining in TS. White vertical tracks indicate the bilateral optical fiber placements in TS. BLA, basolateral amygdala; CEA, central nucleus of the amygdala Scale bar: 0.5 mm. **C)** Top: example histological image showing Cre-dependent expression of ChR2-EYFP (green) along with immunostaining for TH (red) and DAPI (blue) staining in the midbrain. Scale bar: 0.5 mm. Bottom: confocal images showing expression of ChR2-EYFP and TH staining and the merged image showing co- expression. Scale bar: 20 μm. **D)** Schematic of the behavioral protocol. FC: fear conditioning. **E)** Schematic of optogenetic excitation of DA terminals paired to the time of the US (US-Paired) during FC. **F)** Schematic of optogenetic excitation of DA terminals paired to the CS (CS-Paired) during FC. **G)** Percent freezing to the CS during fear conditioning and fear recall sessions. Animals receiving optogenetic excitation at the time of the US (US-Paired ChR2, n = 8) showed enhanced fear learning (two-way repeated measures ANOVA, main effect of group: F_1,56_ = 5.87, p = 0.029; group × trial interaction: F_4,56_ = 3.31, p = 0.016) and fear recall (two-way repeated measures ANOVA, main effect of group: F_1,56_ = 11.57, p = 0.0043) compared to US-Paired EYFP group (n = 8). **H)** Percent freezing to the CS during fear conditioning and fear recall sessions. Animals receiving optogenetic excitation during the CS (CS-Paired ChR2, n = 7) showed enhanced fear learning (two-way repeated measures ANOVA, main effect of group: F_1,44_ = 14.39, p = 0.003; group × trial interaction: F_4,44_ = 3.54, p = 0.013) and fear recall (two-way repeated measures ANOVA, main effect of group: F_1,44_ = 11.66, p = 0.0058) compared to CS-Paired EYFP group (n = 6). **I)** Top: schematic of the real- time place preference test. Bottom: Difference between the percent of time mice spent in laser ON minus laser OFF side for EYFP (n = 8) and ChR2 (n = 8) groups. There was no difference between the two groups (p = 0.56, unpaired t-test), suggesting activation of DA terminals in the TS per se did not induce aversion. **J)** Top: schematic of the open field test. Bottom: percent time mice spent in the center of the open field during laser ON and OFF epochs for EYFP (n = 8) and ChR2 (n = 8) groups. The two groups behaved comparably (two-way repeated measures ANOVA, no main effect of group: F_1,28_ = 0.18, p = 0.67; no group × trial interaction: F_2,28_ = 0.14, p = 0.86). **K)** Top: schematic of the elevated plus maze test. Bottom: percent time mice spent in the open arms of the elevated plus maze during laser ON and OFF epochs for EYFP (n = 8) and ChR2 (n = 8) groups. The two groups behaved comparably (two-way repeated measures ANOVA, no main effect of group: F_1,28_ = 1.88, p = 0.19; no group × trial interaction: F_2,28_ = 0.10, p = 0.90), suggesting that activation of DA terminals in the TS did not cause anxiety-like behaviors. Error bars represent mean ± s.e.m. across animals.

In order to examine enhancement of fear learning, mice were trained in a weak fear conditioning protocol (Figure 5D) using a low US intensity (0.35 mA). The experimental group consisted of ChR2-EYFP expressing mice which received blue light stimulation specifically at the time of the US (US Paired-ChR2, n = 8; Figure 5E). The control group expressing EYFP received the identical light delivery (US Paired-EYFP, n = 8). Comparison of freezing levels to the CS in the two groups revealed a significant difference between the ChR2 and the control mice during both FC and fear recall (Figure 5G). The ChR2 group exhibited higher freezing levels to the CS compared to control mice suggesting enhanced fear conditioning. Thus, excitation of DA terminals in TS at the time of the US is sufficient to enhance associative fear learning.

We next asked whether DA activity in TS during the CS was critical for driving associative fear learning. To address this, we optogenetically excited DA terminals in TS during the CS presentations of FC (Figure 5F). The ChR2- (n = 7) and EYFP- (n = 6) expressing mice underwent the same FC protocol but this time received light stimulation specifically during the CS presentations (CS-paired). Notably, we found a significant difference between the two groups during both FC and fear recall test (Figure 5H). The ChR2-expressing mice froze more to the CS compared to EYFP controls, indicating that excitation of DA terminals in TS during the CS is also sufficient to enhance associative fear learning.

However, excitation of DA neuron terminals in the TS per se could have caused aversion or increased anxiety and hence could have resulted in increased freezing levels rather than enhancing the associative learning process. To examine this possibility, we performed real-time place preference, open field and elevated plus maze tests (Figure 5 I-K) to examine the effect of DA terminal excitation on avoidance and anxiety-like behaviors. Exciting DA terminals in TS (ChR2-EYFP mice n = 8, EYFP mice n = 8) did not cause real-time place avoidance (Figure 5I) suggesting that DA terminal excitation per se was not aversive. Furthermore, we also did not find a significant difference between the two groups in their anxiety-like behaviors when DA terminals in TS were illuminated in the open field (Figure 5J) and the elevated plus maze (Figure 5K). Together, these results suggest that the observed effect on fear learning cannot simply be due to aversion or increased anxiety caused by the excitation of DA terminals in TS.

### Neuronal Activity in TS exhibits a PE-like pattern and potentiation of CS responses during associative fear learning

The requirement of a DA PE signal in TS during FC suggests that activity of TS neurons is likely critical for acquisition of associative fear learning. To address this, we first examined the neuronal activity in TS during FC by measuring activity-dependent Ca^+2^ signals using fiber photometry. To this end, an AAV expressing the Ca^+2^ indicator GCaMP6f was injected and an optical fiber was implanted in the TS (Figure 6A-B, Supplementary Figure 4). The mice (n = 9) underwent the same FC protocol as in fiber photometry experiments described above (Figure 6C), and exhibited successful fear learning (Figure 6D). Interestingly, we observed that the activity in TS exhibited a PE-like pattern during FC (Figure 6E), similar to the results of our recordings of DA terminal activity as well as DA release in TS. While the responses to the US decreased during the course of FC, the CS responses gradually increased (Figure 6E). Indeed, there was a significant increase in responses to the CS from first to last CSs (Figure 6F). Conversely, the responses to the US were larger to the first compared to the last US (Figure 6F). Importantly, we found a significant increase in responses to the CS from Hab to fear recall (Figure 6G). In almost all recording sites, the CS responses during fear recall were larger in amplitude compared to Hab (Figure 6G). Together, these results demonstrated that a PE-like activity pattern was observed in the TS during associative fear learning.

**Figure 6.**
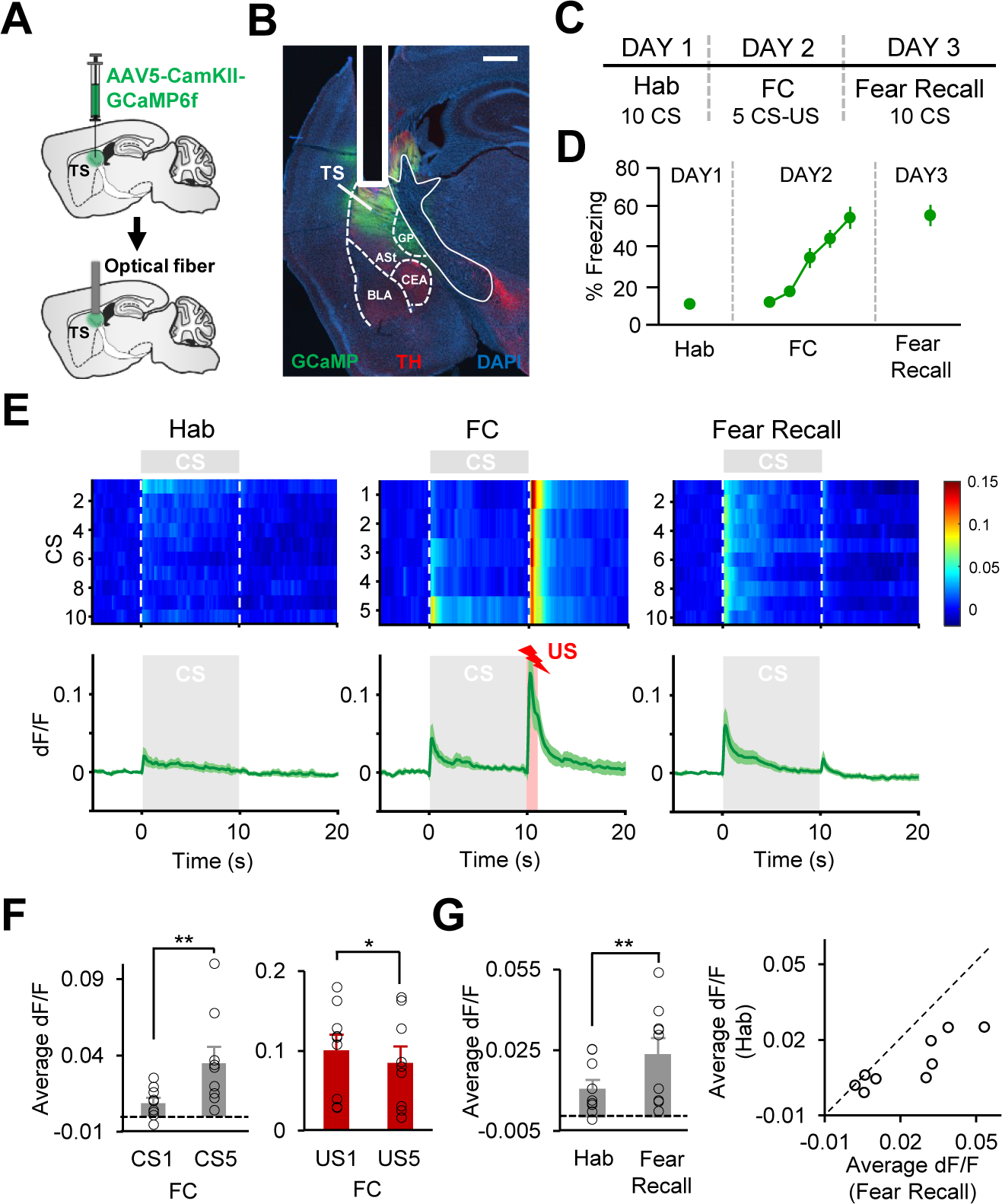
Activity in TS exhibits a PE-like pattern and CS responses are potentiated during associative fear learning. A) Schematic of the surgical procedure showing virus injection (top) and optical fiber implantation (bottom) in TS. **B)** Example histological image showing expression of GCaMP (green) along with immunostaining for tyrosine hydroxylase (TH, red) and DAPI (blue) staining in TS. White vertical track indicates the optical fiber placement in TS. ASt, amygdalostriatal transition area; BLA, basolateral amygdala; CEA, central nucleus of the amygdala; GP, globus pallidus. Scale bar: 0.5 mm. **C)** Schematic of the behavioral protocol. Hab: tone habituation, FC: fear conditioning. **D)** Behavioral freezing to the CS (n = 9 mice) during Hab, FC and Fear Recall. **E)** Top: Average GCaMP activity around each CS during Hab, FC and Fear Recall across all recording sites (n = 9). The heat map shows response amplitudes (dF/F) around each CS presentation. Bottom: Average change in fluorescence around the time of CS presentation (gray area) during Hab, FC and Fear Recall. The red area during FC represents the US presentation. **F)** Left: comparison of average change in fluorescence in the 5s after CS onset during CS1 and CS5 of FC. Note the significant increase in the Ca^2+^ signal from CS1 to CS5 (**p = 0.0039, signed-rank test). Right: comparison of average change in fluorescence during the US (1s) for US1 and US5 of FC. Note the significant decrease in the Ca^2+^ signal from US1 to US5 (*p = 0.02, signed- rank test). **G)** Left: Average change in fluorescence in the 5s after CS onset during Hab and Fear Recall. Note the significant increase in the Ca^2+^ signal from Hab to Fear Recall (**p = 0.0078, signed-rank test). Right: scatter plot showing CS responses of each recording site (n = 9) during Hab and Fear Recall. Data points below the unity line represent larger CS responses during Fear Recall. Shaded regions and error bars represent mean ± s.e.m across animals.

### Neuronal Activity in TS is required for and sufficient to enhance associative fear learning

We next asked whether activity in TS was necessary for associative fear learning. To address this, we performed chemogenetic inhibition of TS neurons during FC. Mice received injections of an AAV expressing the inhibitory DREADD receptor (hM4D(Gi)) in TS (Figure 7A, B) and underwent the same auditory FC protocol as in the 6-OHDA experiment (Figure 7C). Thirty minutes before the FC session, mice received systemic injections of the DREADD agonist clozapine N-oxide (CNO) to inhibit activity of TS neurons during fear conditioning whereas control mice received saline injections. We found that CNO-injected mice (n = 8) froze significantly less to the CS at the end of FC compared to saline-injected controls (n = 8; Figure 7D, E) suggesting impaired fear learning. Furthermore, impaired fear learning resulted in a weaker fear memory when tested the next day during the fear recall test (Figure 7D, F). These results demonstrated that the activity of TS neurons is critical for associative fear learning.

**Figure 7.**
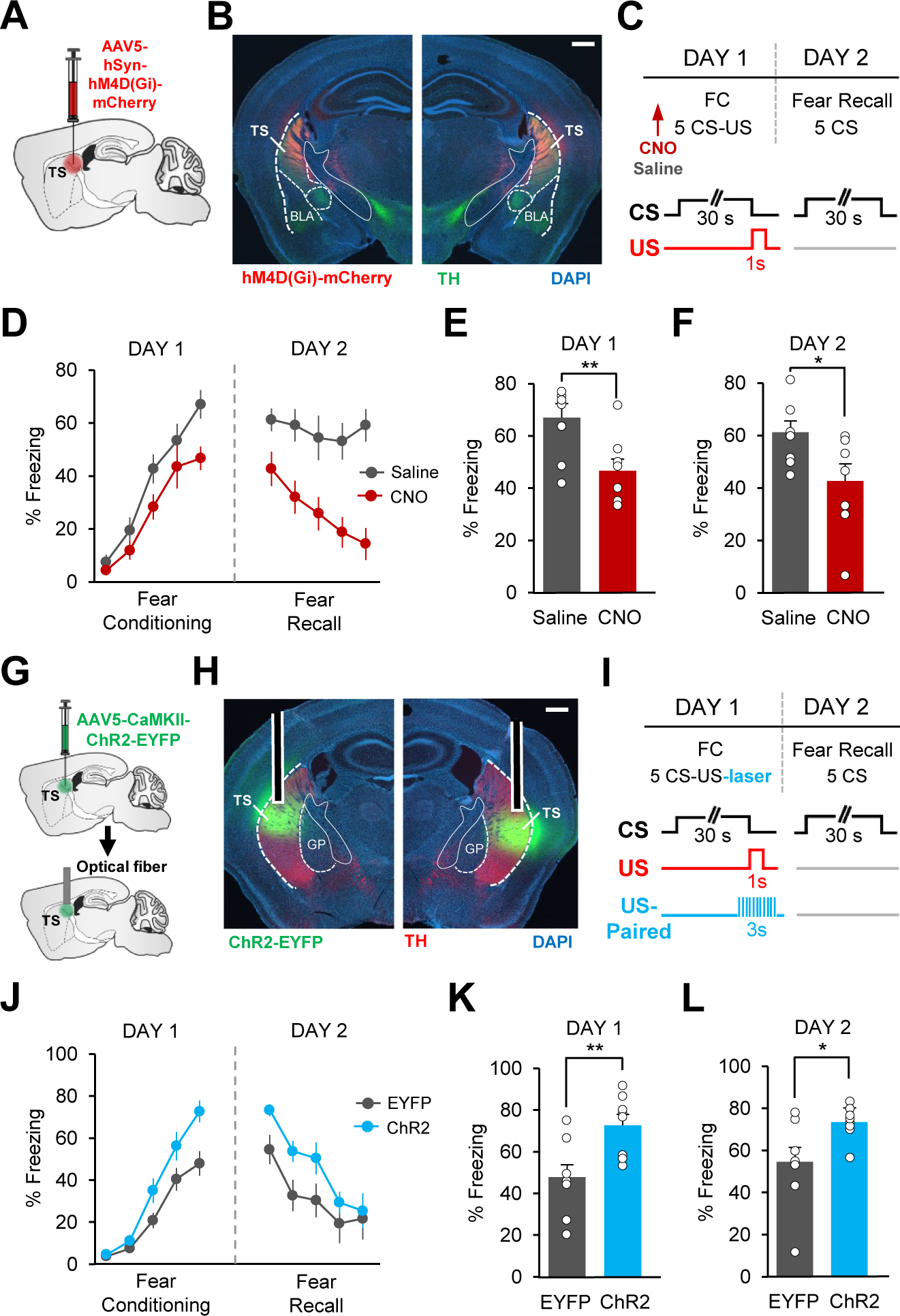
Activity in TS is required for and sufficient to enhance associative fear learning. A) Schematic of the surgical procedure showing virus injection in TS. **B)** Example histological image showing expression of hM4D(Gi)-mCherry (red) along with immunostaining for tyrosine hydroxylase TH (green) and DAPI (blue) staining in TS. Scale bar: 0.5 mm. **C)** Top: schematic of the behavioral protocol. FC: fear conditioning. Bottom: schematic of CS and US presentations during FC and Fear Recall sessions. **D)** Percent freezing to the CS during fear conditioning and fear recall sessions for control saline- (gray, n = 8) and experimental CNO-injected mice (red, n = 8). There was a significant difference between the two groups during fear conditioning (two-way repeated measures ANOVA, main effect of group, F_1,56_ = 5.08, p = 0.04) and fear recall sessions (two-way repeated measures ANOVA, main effect of group, F_1,56_ = 19.61, p = 0.0006), indicating impaired fear learning in the CNO-injected mice. **E)** Freezing to the last CS during fear conditioning. CNO group froze significantly less compared to control mice (t-test, t(14) = 3.22, p = 0.006). **F)** Freezing to the first CS during fear recall. CNO group froze significantly less compared to control mice (t-test, t(14) = 2.38, p = 0.03). **G)** Schematic of the surgical procedure showing virus injection and optical fiber implantation in the TS. **H)** Example histological image showing expression of ChR2-EYFP (green) along with immunostaining for TH (red) and DAPI (blue) staining in the TS. GP, globus pallidus. Scale bar: 0.5 mm. **I)** Top: schematic of the behavioral protocol. Bottom: schematic of CS and US presentations and optogenetic excitation paired to the time of the US (US-Paired) during fear conditioning. **J)** Percent freezing to the CS during fear conditioning and fear recall sessions for control EYFP (gray, n = 8) and experimental ChR2 (blue, n = 8) groups. There was a significant difference between the two groups during fear conditioning (two-way repeated measures ANOVA, main effect of group, F_1,56_ = 5.8, p = 0.03 and group × trial interaction, F_4,56_ = 4.05, p = 0.0059), indicating enhanced fear learning in the ChR2 group. **K)** Freezing to the last CS during fear conditioning. ChR2 group froze significantly more compared to control mice (t-test, t(14) = 3.06, p = 0.008). **L)** Freezing to the first CS during fear recall. ChR2 group froze significantly more compared to EYFP group (t-test, t(14) = 2.3, p = 0.03). Error bars represent mean ± s.e.m. across animals.

A PE-like activity pattern in TS for the aversive US suggests that this signal might drive fear learning. If that is the case, boosting activity of TS neurons during the aversive US is expected to enhance fear learning. We tested this by performing temporally-precise optogenetic excitation of TS neurons at the time of the US during FC (Figure 7I). To this end, an AAV expressing ChR2-EYFP or EYFP only was bilaterally injected and optic fibers were bilaterally implanted in TS (Figure 7G-H, Supplementary Figure 5). In order to examine enhancement of fear learning, mice were trained in a weak fear conditioning protocol (Figure 7I), as during optogenetic excitation of DA terminals described above (Figure 5). We found a significant difference between the ChR2 (n =8) and the control (n = 8) mice during FC (Figure 7J). ChR2-expressing mice showed significantly higher freezing to the CS at the end of FC (Figure 7K) and the beginning of fear recall test (Figure 7L), suggesting enhanced fear learning. Together, these results demonstrate that excitation of TS neurons at the time of the US is sufficient to enhance associative fear learning.

### DA input is critical for PE-like activity and potentiation of CS responses in TS during associative fear learning

Our results demonstrated that the activity in TS exhibits a PE-like pattern during fear learning. A key question is whether DA input is crucial for this fear learning-related activity observed in TS. To address this question, we performed 6-OHDA ablation of TS-projecting DA neurons followed by Ca^2+^ recordings in TS using fiber photometry. Mice received 6-OHDA or saline injections, and one week later, were injected with an AAV expressing GCaMP6f as well as implanted with an optical fiber in the TS (Figure 8A, Supplementary Figure 6). The animals underwent the same FC protocol (Figure 8B) as in fiber photometry experiments described above. Importantly, 6-OHDA injected animals (n = 8) showed reduction in DAergic innervation of TS compared to control group (n = 7; Figure 8D, H), and consistently, lesioned mice showed impaired fear learning compared to controls (Figure 8C).

**Figure 8.**
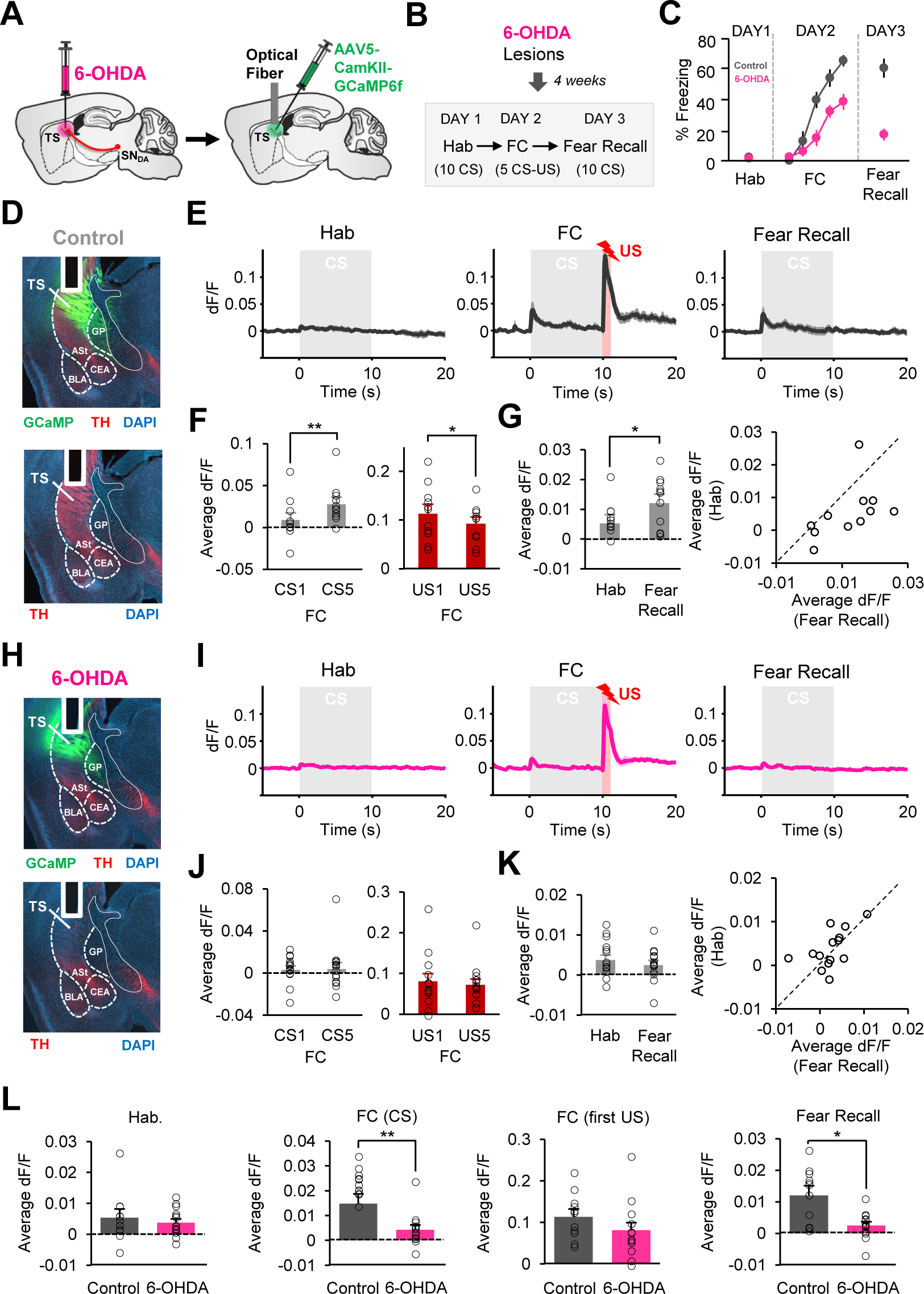
DA input is critical for PE-like activity and potentiation of CS responses in TS during associative fear learning. A) Schematic of the surgical procedure. **B)** Schematic of the behavioral procedure. Hab: tone habituation, FC: fear conditioning. **C)** Behavioral freezing to the CS during Hab, FC and fear recall in control (n = 7) and 6-OHDA (n = 8) groups. **D, H)** Example histological image showing expression of GCaMP (green; top) along with immunostaining for tyrosine hydroxylase (TH, red) and DAPI (blue) staining in TS of a control **(D)** and a 6-OHDA lesioned **(H)** animal. Note the decrease in TH staining in the TS of 6-OHDA injected animals indicating ablation of TS-projecting DA neurons. White vertical track indicates the optical fiber placement in TS. ASt, amygdalostriatal transition area; BLA, basolateral amygdala; CEA, central nucleus of the amygdala; GP, globus pallidus. **E, I)** Average change in fluorescence around the time of CS presentation (gray area) during Hab, FC and Fear Recall in control (**E,** n = 11 recording sites) and 6-OHDA (**I,** n = 14 recording sites) groups. **F, J)** Left: average change in fluorescence in the 5s after CS onset during CS1 and CS5 of FC in the control **(F)** and 6-OHDA **(J)** groups. Note the significant increase in the Ca^2+^ signal from CS1 to CS5 in the control (**p = 0.0049, signed-rank test) but not 6-OHDA lesioned (p = 0.5, signed-rank test) mice. Right: average change in fluorescence during the US (1s) for US1 and US5 of FC. Note the significant decrease in the Ca^2+^ signal from US1 to US5 in the control (*p = 0.024, signed-rank test) but not 6-OHDA lesioned (p = 0.62, signed-rank test) mice. **G, K)** Left: average change in fluorescence in the 5s after CS onset during Hab and Fear Recall in the control **(G)** and 6-OHDA lesioned **(K)** mice. The control, but not 6- OHDA, group exhibited a significant increase in the Ca^2+^ signal from Hab to Fear Recall (control: *p = 0.036; 6-OHDA: p = 0.23, signed-rank test). Right: Scatter plots showing CS responses of each recording site in the control **(G)** and 6-OHDA **(K)** mice during Hab and Fear Recall. Data points below the unity line represent larger CS responses during Fear Recall. **L)** Comparison of the average Ca^2+^ signal in response to the CS during Hab (left; p = 0.80, rank-sum test) and Fear Recall (right; p = 0.014, rank-sum test), and CS (average of 5 CSs; p = 0.006, rank-sum test) and unpredicted US (first US) during FC (middle; p = 0.13, rank-sum test) between control and 6- OHDA groups. Note the significant difference between the two groups in their CS responses during FC and Fear Recall indicating enhancement of CS responses as a result of associative learning in the control, but not the 6-OHDA, group. Shaded regions and error bars represent mean ± s.e.m across animals.

The PE-like activity pattern during FC that the control group exhibited (n = 11 recording sites; Figure 8E-G) was not observed in the 6-OHDA lesioned group (n = 14 recording sites; Figure 8I-K). In lesioned mice, we did not find an increase in the CS responses from first to last CSs (Figure 8J), which was seen in control animals (Figure 8F). Interestingly, the significant decrease in responses between the first and the last USs of FC, that was observed in control mice (Figure 8F), was also absent in lesioned animals (Figure 8J). Furthermore, while CS responses were significantly increased between Hab and fear recall in control animals, no increase was observed in lesioned mice (Figure 8G, K). Whereas CS responses were larger in magnitude during fear recall compared to Hab in the majority of control animals, this was not the case in lesioned mice (Figure 8G, K). Notably, there was no difference between the two groups in their CS responses during Hab, whereas during FC and fear recall, CS responses were significantly larger in the control group (Figure 8L). Importantly, the responses to unpredicted US (first US of FC) were not significantly different between the two groups (Figure 8L), suggesting that US input to TS in general was not affected. Rather, our results demonstrate that the decrease in US responses as fear conditioning progressed was absent, consistent with a deficit in learning about the CS-US association in 6-OHDA lesioned mice. Overall, these results indicate that DA input is crucial for fear learning-induced activity patterns observed in TS during associative fear learning.

## DISCUSSION

Whether midbrain DA neurons projecting to brain structures outside the canonical amygdala circuitry encode an aversive PE signal that is crucial to drive associative fear learning is largely unknown. Recent studies have implicated TS in fear learning (Kintscher et al., 2023; Chen et al., 2023), yet the precise role DA neurons projecting to TS play in associative fear learning is incompletely understood. Here, we demonstrated that DA projections to TS drive associative fear learning by encoding an aversive PE signal which is crucial for generating fear learning-induced activity patterns in TS. We first showed that DA projections to TS exhibit a positive PE signal during FC, and that this PE signal is transmitted by DA release in TS. Projection-specific lesioning of TS-projecting DA neurons selectively impaired the acquisition of associative fear learning, but not fear retrieval and expression. Notably, these neurons were required specifically for acquiring the association between the auditory CS and US, but not between the context and the US or between an auditory CS and reward. Temporally-precise optogenetic excitation of DA projections to TS enhanced associative fear learning. Furthermore, activity in TS exhibited a PE-like pattern during FC, and the neuronal activity in TS was required for and sufficient to enhance fear learning. Finally, we demonstrated that DA input was crucial for the fear learning-induced activity pattern in TS.

The midbrain DA system is composed of functionally distinct and mostly non-overlapping subpopulations of DA neurons, each of which projects mainly to a single brain region (Beier et al., 2015; Farassat et al., 2019; Lammel et al., 2008; Lerner, et al., 2015; Menegas et al., 2015; Roeper, 2013). Notably, DA neurons projecting to TS have recently emerged as a unique subpopulation based on their distinct input-output circuitry (Menegas et al., 2015). These DA neurons have been shown to be activated by novel as well as a subset of aversive stimuli (e.g. air puffs and loud tones but not bitter taste; Menegas et al., 2017; 2018). In line with these previous studies, we found that TS-projecting DA neurons are activated particularly strongly by aversive stimuli that are noxious such as foot shocks, but only weakly by rewards. Interestingly, TS-projecting DA neurons have been shown to exhibit a PE-like activity pattern during Pavlovian association of olfactory cues with mildly aversive stimuli such as air puffs (Menegas et al., 2017; 2018); and these DA neurons are particularly important for novelty-induced threat avoidance (Akiti et al., 2022; Menegas et al., 2018). However, whether TS-projecting DA neurons encode a PE signal for strong painful stimuli such as foot shocks; and whether this signal is necessary for associating cues with the aversive US during FC has remained elusive.

During associative learning, a positive PE acts as a teaching signal and drives learning about the CS that precedes the US (Rescorla and Wagner, 1972; Glimcher, 2011; Montague et al., 1996; Schultz and Dickinson, 2000; Steinberg et al., 2013). We here demonstrate that TS-projecting DA neurons encode a PE for the aversive US during FC. These neurons respond more strongly to aversive USs when they are unpredicted compared to predicted ones. Furthermore, we observed that during the course of FC the US responses decreased whereas the CS responses were enhanced as the CS-US association was learned and the CS came to predict the US. These changes in the activity of TS-projecting DA neurons were specifically dependent on the temporally contingent presentation of the CS and the US: in mice trained with unpaired conditioning, during which the CS and the US were presented in a temporally unpaired manner and thus CS did not predict the occurrence of the US, the CS responses failed to potentiate. Notably, a PE signal is expected to initiate new learning; consistent with this, we demonstrate that TS-projecting DA neurons are required selectively during the acquisition of the CS-US association during FC, but not for retrieval and expression of fear memories. Interestingly, ablation of TS-projecting DA neurons did not have an effect on acquisition of contextual fear memories, highlighting the selective role of these DA neurons in cued associative fear learning. Furthermore, we demonstrate that exciting DA projections to TS at the time of the US, which boosts the aversive PE signal, caused enhancement of FC. Together, our results indicate that the aversive PE signal encoded by TS-projecting DA neurons indeed acts as a teaching signal to drive the acquisition of CS-US association during fear learning.

In contrast to their strong responses to the aversive US, we show that DA projections to TS are only weakly activated by rewards. Consistent with this, we found that TS-projecting DA neurons were not required for associating an auditory cue with reward; although they have been implicated in more complex forms of reward learning involving auditory discrimination (Chen et al., 2022). In line with our findings, TS-projecting DA neurons do not exhibit RPE signaling during Pavlovian reward learning tasks (Menegas et al., 2017). Overall, our results thus indicate an aversive bias in the responses of TS-projecting DA neurons. This distinguishes the responses of TS-projecting DA neurons from other subpopulations of DA neurons that project to the striatum, in particular the ones projecting to lateral and ventral subregions of NAc which have also been shown to be activated by foot shocks (de Jong et al., 2019; Salinas-Hernández, 2023; Yuan et al., 2019). However, those DA neurons also exhibit strong, and even larger, responses to rewards compared to aversive stimuli (de Jong et al., 2019; Salinas-Hernández, 2023), suggesting that they likely encode salience rather than aversion. Our results raise the question of whether activation of TS-projecting DA neurons induces aversion per se (Verharen et al., 2020; Zafiri and Duvarci, 2022). However, we found that exciting DA terminals in TS does not cause place avoidance or anxiety suggesting that this DA input is likely not aversive by itself. Instead, our results suggest that DA input likely induces plasticity in TS that biases associative learning to cues that are specifically paired with aversive outcomes.

The amygdala, in particular LA, is established to be the critical site for plasticity mediating the CS-US association during fear learning (Duvarci and Paré, 2014; Johansen et al., 2011; LeDoux, 2000; Maren and Quirk, 2004; Pape and Pare, 2010; Tovote et al., 2015). LA receives inputs from both the thalamus and the cortex, relaying information about the auditory CS (Romanski et al., 1993). Considerable evidence indicates that potentiation of CS-evoked responses in LA neurons underlies acquisition of associative fear learning (Quirk et al., 1995; Rogan et al., 1997; Collins and Paré, 2000; Repa et al., 2001; Rosenkranz and Grace, 2002; Goosens et al., 2003). Moreover, plastic changes in the activity of neurons located in the central nucleus of the amygdala (CEA) have also been shown to underlie acquisition of fear memories (Ciocchi et al., 2010; Duvarci et al, 2011; Haubensak et al., 2010; Wilensky et al., 2006). In line with these findings, DA projections to the amygdala exhibit activation during fear learning (Groessl et al., 2018; Lutas et al., 2019; Tang et al., 2020); and in particular, non-canonical dorsal tegmental DA neurons, that are located in the periaqueductal gray (PAG)/dorsal raphe (DR) and project specifically to CEA, have been shown to encode a PE signal to gate fear learning (Groessl et al., 2018). However, whether DA signaling outside the amygdala circuitry contributes to the acquisition of CS-US association during FC has remained largely unknown. Our findings reveal, for the first time, that DA neurons projecting to a brain structure outside the canonical amygdala circuitry, encode an aversive PE signal crucial for driving associative fear learning.

While the role of the amygdala is well-established, it is becoming increasingly clear that acquisition of fear memories also involves plasticity in brain structures beyond the traditional amygdala circuitry (Herry and Johansen, 2014; Dalmay et al., 2019; Letzkus et al., 2011; Weinberger, 2011). Notably, despite earlier lesioning and anatomical studies hinting at the involvement of the posterior striatal areas, including the TS, in fear conditioning (LeDoux et al., 1985; 1986; 1991), the contribution of TS in fear learning remained largely elusive and has recently begun to be investigated (Kintscher et al., 2023; Chen et al., 2023). TS, also known as the auditory striatum, is a distinct subregion of the dorsal striatum, characterized by the unique set of inputs it receives (Hunnicutt et al., 2016, Valjent and Gangarossa, 2021). Similar to LA, TS receives auditory inputs from both the thalamus and the cortex (LeDoux et al., 1985; 1991); and we here showed that auditory CS-evoked responses are potentiated also in the TS as the animals learned the CS-US association during FC, consistent with previous reports (Kintscher et al., 2023; Chen et al., 2023). Importantly, we demonstrate that neuronal activity in TS is crucial for acquisition of fear conditioning, indicating that a broader neural network than the amygdala circuitry is indeed involved in acquiring the CS-US association during fear learning.

A striking finding in our study is the PE-like activity pattern that we observed in TS. Similar to TS-projecting DA neurons, TS neuronal activity exhibited larger responses to the US at the beginning of FC when US presentation was unpredicted. As the CS-US association was learned, the US responses decreased, and conversely, responses to the CS gradually increased. Importantly, we found that PE coding during FC was dependent on the DA input to TS. We found that the US responses did not decrease during the course of FC in mice with ablation of TS-projection DA neurons, consistent with the deficit in acquiring the CS-US association in these mice. Notably, responses to the first unpredicted US were not different between the lesioned and control mice indicating that the somatosensory inputs to TS relaying the US information were not affected by ablation of the DA input, but rather the predictive coding of US responses in TS was impaired. Furthermore, CS responses did not potentiate in parallel with the impaired fear learning in lesioned mice. Together, these results demonstrate that the fear learning-induced activity pattern in TS required DA input during FC. PE signaling during FC has previously been demonstrated in LA, where the PE coding was shown to set the associative memory strength (Ozawa et al., 2017). In line with this notion, we here demonstrated that optogenetic excitation of TS neuronal activity at the time of the US enhanced fear learning, indicating that the aversive PE coding by TS neurons drives learning about the CS-US association. Aversive PE coding in LA was shown to be mediated by a feedback neuronal circuitry involving projections form ventrolateral PAG (vlPAG) to LA (Johansen et al., 2010; Ozawa et al., 2017), and likely also involves projections from the cerebellum to vlPAG (Frontera et al., 2020; Urrutia Desmaison et al., 2023). Whether feedback neural circuits are recruited to drive aversive PE coding in TS neurons, as well as in TS-projecting DA neurons, and the components of these neural circuits will be important questions for future research. Another important question that remains to be investigated is the differential contributions of LA versus TS neuronal activity in driving associative fear learning. Overall, our study, for the first time, reveals that a DA PE signal in a non-canonical nigrostriatal circuitry is crucial for driving associative fear learning.

## METHODS

### Subjects

All procedures were conducted in accordance with the guidelines of the German Animal Welfare Act and were approved by the local authorities (Regierungspräsidium Darmstadt). Adult male C57BL/6N (Charles River or Janvier Labs) and heterozygous DAT-Cre mice (Zhuang et al. 2005; backcrossed with C57BL/6N) aged older than 3 months at the start of experiments were used. All experimental groups were matched for age. For 6-OHDA lesioning, optogenetic and chemogenetic experiments, littermate mice were allocated to experimental and control groups.

All mice were individually housed on a 12-h light/dark cycle. All experiments were performed during the light cycle.

### Viral Constructs

We obtained AAV5-CAG-Flex-GCaMP6m-WPRE-SV40, pENN-AAV5-CaMKII-GCaMP6f-WPRE-SV40, AAV5-CAG-dLight1.1 and AAV5-hSyn-hM4D(Gi)-mCherry from Addgene, and AAV5-EF1a-DIO-hChR2(H134R)-EYFP, AAV5-EF1a-DIO-EYFP, AAV5-CaMKIIa-hChR2(H134R)-EYFP, AAV5-CaMKIIa-EYFP and AAV5-CaMKII-GFP from the University of North Carolina Vector Core.

### Surgical procedures

Animals were anesthetized using isoflurane (1–2%) and placed in a stereotaxic frame. At the onset of anesthesia, all animals received intraperitoneal injections of atropine (0.05 mg/kg) and subcutaneous injections of carprofen (4 mg/kg) and dexamethasone (2 mg/kg). Eye gel was applied on the eyes to prevent dehydration of the cornea. Lidocaine cream was applied on the scalp as local anesthetic. The animal’s temperature was maintained for the duration of the surgical procedure using a heating blanket. Anesthesia levels were monitored throughout the surgery and the concentration of isoflurane adjusted so that the breathing rate never fell below 1 Hz.

For GCaMP fiber photometry recordings of DA terminal activity in the TS, DAT-cre mice were injected with 0.5-1 μl of AAV5-CAG-Flex-GCaMP6m-WPRE-SV40 (final titer ∼1 x 10^13^ pp per ml) in the SN (3.2 mm posterior to bregma, 1.2 mm lateral to the midline and 4.5 mm ventral to bregma) at 50 nl/min using a 10 μl syringe with a 33-gauge needle controlled by an injection pump. The needle was left in place for an additional 10-15 min before slowly being withdrawn.

Following infusion of the virus, optical fibers (400 μm core diameter, 0.48 NA, Doric Lenses) were slowly inserted through the craniotomy above the TS (AP: -1.3 mm, ML: 2.95 mm and DV: 2.75 – 3.0 mm). In a subset of animals, recordings were performed bilaterally. The optical fiber was then anchored to the skull using skull screws and dental cement (Paladur).

For GCaMP fiber photometry recordings of neuronal activity in the TS, wild-type C57BL/6N mice were injected with 150 nl of AAV5-CaMKII-GCaMP6f-WPRE-SV40 (final titer ∼1 x 10^13^ pp per ml) in the TS (AP: -1.3 mm, ML: 2.95 mm and DV: 3.25 mm) at 50 nl/min using a 10 μl syringe with a 33-gauge needle controlled by an injection pump. The needle was left in place for an additional 10-15 min before slowly being withdrawn. Following infusion of the virus, optical fibers (400 μm core diameter, 0.48 NA, Doric Lenses) were slowly inserted through the same craniotomy to a depth of 2.75 – 3.0 mm below the bregma. The optical fiber was then anchored to the skull using skull screws and dental cement (Paladur).

For dLight fiber photometry experiments, wild-type C57BL/6N mice were injected with 150-300 nl of AAV5-CAG-dLight1.1 (final titer 4.2 x 10^12^ pp per ml) in the TS (AP: -1.3 mm, ML: 2.95 mm and DV: 3.25 mm) as described above. Following infusion of the virus, an optical fiber (400 μm core diameter, 0.48 NA, Doric Lenses) was slowly inserted through the same craniotomy to a depth of 2.75 – 3.0 mm below the bregma. In a subset of animals, recordings were performed bilaterally. The optical fibers were then anchored to the skull using skull screws and dental cement (Paladur).

6-OHDA lesioning of TS-projecting DA neurons was performed as previously described (Menegas et al., 2018; Thiele et al. 2012). Thirty min before the surgery, wild-type C57BL/6N mice received intraperitoneal injections of a solution (10ml/kg) containing desipramine hydrochloride (2.5 mg/ml, Sigma-Aldrich) and pargyline hydrochloride (0.5 mg/ml, Sigma-Aldrich) dissolved in 0.9% saline (pH 7.4), in order to prevent uptake of 6-OHDA in noradrenaline neurons. 6-OHDA (10mg/ml; 6-OHDA hydrobromide, Sigma-Aldrich) was dissolved in a vehicle solution immediately before the surgeries. The vehicle contained saline (0.9%) and a small amount of ascorbic acid (0.2%) to prevent oxidation of 6-OHDA. To further minimize oxidation of 6-OHDA, the solution was kept on ice, wrapped in aluminum foil and used within 4h after preparation. During the surgery, animals were injected bilaterally with 150-300 nl of the 6-OHDA solution or the vehicle in the TS using the coordinates described above. The injections were performed at 50 nl/min speed using a 10 μl syringe with a 33-gauge needle which was controlled by an injection pump. The needle was left in place for an additional 10-15 min before slowly being withdrawn. The scalp incision was then closed using sutures. In experiments with GCaMP recordings, mice received virus injections and optical fiber implantation as described above one week after 6-OHDA injections. In a subset of animals in the control and 6-OHDA groups, recordings were performed bilaterally.

For optogenetic excitation of DA terminals in the TS, DAT-cre mice were injected bilaterally in the SN with 0.5-1 μl of AAV5-EF1a-DIO-hChR2(H134R)-EYFP (final titer 4.4 x 10^12^ pp per ml) or AAV5-EF1a-DIO-EYFP (final titer 4.3 x 10^12^ pp per ml) per hemisphere using the coordinates described above. Virus injection was performed as described above and followed by implantation of optical fibers (200 μm core diameter, 0.22 NA, Thorlabs) bilaterally above the TS (AP: -1.3 mm, ML: 2.95 mm) to a depth of 2.75 – 3.0 mm below bregma. The optical fibers were then anchored to the skull using skull screws and dental cement (Paladur).

For optogenetic excitation of neuronal activity in the TS, wild-type C57BL/6N mice were injected bilaterally in the TS with 150 nl of AAV5-CaMKIIa-hChR2(H134R)-EYFP (final titer 4.1 x 10^12^ pp per ml) or AAV5-CaMKIIa-EYFP (final titer 3.6 x 10^12^ pp per ml) per hemisphere using the coordinates described above. Virus injection was performed as described above. Optical fibers (200 μm core diameter, 0.22 NA, Thorlabs) were then implanted bilaterally above the TS using the coordinates described above. The optical fibers were then anchored to the skull using skull screws and dental cement (Paladur).

For chemogenetic experiments, C57BL/6N mice were bilaterally injected with 150 nl of AAV5- hSyn-hM4D(Gi)-mCherry (final titer 8.6 x 10^12^ particles per ml) per each hemisphere in the TS using the coordinates described above. Virus injection was performed as described above. The scalp incision was then closed using sutures at the end of the surgery.

### Behavior

*Fear conditioning*. Fear conditioning and fear recall took place in two different contexts (A and B). Context A consisted of a square chamber with an electrical grid floor (Med Associates) used to deliver the footshock US. Context B consisted of a white teflon cylindrical chamber with bedding material on the floor. The chambers were located inside a sound attenuating box and were cleaned with 1% acetic acid before and after each session. Before fear conditioning experiments started, all mice were habituated to handling and being connected to the patch cord and habituated to contexts A and B for 10-15 min each in a counterbalanced fashion. On day 1, mice received a tone habituation session which started following a 2 min baseline period in context A and consisted of 10 presentations of the CS (4 kHz tone, 75 dB) with a random intertrial interval (ITI) of 40-120 s. On day 2, mice underwent fear conditioning consisting of five pairings of the CS with a US (1s footshock, ITI: 40-120 s). The CS was 10s and 30s long in photometry and optogenetic/chemogenetic experiments, respectively. In 6-OHDA experiment, both 10s and 30s long CSs were used. Comparable results were obtained with both CS durations (p > 0.05) and hence the data was pooled. The US intensity was 0.4 – 0.5 mA for photometry and 0.5 mA for 6-OHDA and chemogenetic experiments. For optogenetic excitation experiments where mice received a weak training, the US intensity was 0.35 mA. The onset of the US coincided with the offset of the CS. On day 3, mice received a fear recall session consisting of 10 presentations of the CS alone in context B for photometry experiments. For 6- OHDA, optogenetic and chemogenetic experiments, fear recall test consisted of 5 presentations of the CS on day 3. For contextual fear conditioning, mice received context habituation (context A) for 10 min on day 1. On day 2, following a 2 min baseline period mice received 5 presentations of the US (0.5 mA) with a random intertrial interval (ITI) of 40-120 s in context A. On day 3, mice were placed back to context A for 10 min for their context testing. For the unpaired training protocol, mice received five presentations of the CS and the US in an explicitly unpaired fashion (ITI: 40-120 s) on day 2. At the end of fiber photometry experiments, a subset of mice was further tested for predicted versus unpredicted US presentations (5 CS-US pairings and 5 USs presented in a random order). Throughout the experiments, the behavior of mice was recorded to video and scored by experienced observers blind to the experimental condition. Behavioral freezing, defined as the absence of all bodily movements except breathing-related movement (Blanchard and Blanchard, 1972), was used as the measure of fear.

*Reward Tasks.* Mice in the photometry experiments underwent a reward task following the fear conditioning protocol. To motivate animals, their access to water was restricted until they reached 85% of their body weight. Animals were placed in an operant chamber containing a liquid delivery port. Nosepokes into the delivery port that followed the previous nosepoke by at least a variable inter-trial interval of 3-5s triggered the delivery of liquid reward (10% sucrose solution). Animals were first trained on 90% probability of reward delivery and the next day photometry recordings were performed with a 50% reward probability.

In 6-OHDA experiment, mice underwent a Pavlovian reward learning task for 5 consecutive days. Animals were placed in an operant chamber containing a liquid delivery port. Each session consisted of 50 pairings of a CS (8kHz tone, 75 dB, 5s long) and a reward (10% sucrose solution) with a random ITI of 20-30 s. If the animal accessed the reward port during the CS presentation, a 10 μl reward was delivered.

*Real-time place preference test*. At the end of fear conditioning protocol, a subset of mice in the optogenetic experiments underwent place preference, open field and elevated plus maze tests. Real-time place preference test was conducted in a custom-made chamber (50 x 50 x 50 cm, wooden gray box) divided into two compartments. The test consisted of two 10 min phases (Figure 5I). During the first phase, one side of the chamber was randomly assigned as the laser ON side. Mice were individually connected to the patch cords and placed in the laser OFF side of the chamber. Each time the mouse entered the laser ON side laser light was delivered until the mouse crossed back to the OFF side. In the second phase, the sides were switched and the previously laser OFF side became laser ON side in order to counterbalance each side.

*Open field test*. The custom made open field chamber (50 x 50 x 50 cm, wooden gray box) was divided into a central area (center 25 x 25 cm) and an outer area (periphery). The open field test consisted of a 9 minute session with three alternating 3-minute epochs (OFF-ON-OFF epochs) in which laser was delivered during the ON epoch (Figure 5J).

*Elevated Plus Maze (EPM)*. EPM consisted of two open arms (30 × 5 cm), two closed arms (30 × 5 × 15 cm) and a central area (5 × 5 × 5 cm). The maze was placed 40 cm above the floor. Mice were individually connected to the patch cords and placed in the center. The test consisted of a 9 minute session with three alternating 3-minute epochs (OFF-ON-OFF epochs) in which laser was delivered during the ON epoch (Figure 5K).

### GCaMP and dLight recordings using fiber photometry

Animals were injected with viral vectors and implanted with optical fibers in the TS, as described above. After a waiting period of 3-4 weeks to allow for surgical recovery and virus expression, mice were connected to 400 μm patch cords (Doric Lenses). Fluorescence was measured by delivering 465 nm excitation light through the patch cord and separating the emission light at 525 nm with a beamsplitter (Fluorescence MiniCube FMC3, Doric Lenses). The emission light was collected using a Femtowatt Silicon Photoreceiver (Model # 2151, Newport). The voltage output of the photoreceiver was then digitized at 2 kHz (Digital Lynx SX, Neuralynx). After animals were habituated to handling and being connected to the patch cord, they underwent the behavioral protocol as described above. Photometry recordings were performed throughout the protocol.

### Analysis of fiber photometry data

The voltage output of the photoreceiver, representing fluctuations in fluorescence, was downsampled to 10 Hz. The change in fluorescence evoked by the CS (dF/F) was then calculated by subtracting from each trace the baseline fluorescence (average during the 5s before CS onset) and dividing it by the baseline fluorescence. dF/F traces were then averaged separately for each animal for tone habituation (Hab: average of 10 CSs), fear conditioning (Fear Cond., average of 5 CSs) and fear recall (average of 10 CSs). To examine responses to the CS, we further averaged dF/F values in the 5s following CS onset in each session. To quantify responses to the US, we averaged the dF/F values at the time of the US (1s) during Fear Cond. for each animal. To examine the change in responses to the CS and the US during Fear Cond., we compared responses to the first and last CSs and USs for each animal. To examine responses to reward, average dF/F was calculated for rewarded trials using the baseline fluorescence 3s before noseport entry. Reward responses were quantified by averaging the dF/F in the 3s following noise port entry for each animal.

### Chemogenetic experiments

Three to four weeks after viral injections, mice underwent the chemogenetic inhibition experiment. Thirty min before the fear conditioning session, mice received systemic injections of the DREADD agonist clozapine N-oxide (CNO, Sigma-Aldrich; 1-1.5 mg/kg dissolved in saline). The control group was injected with saline. Fear conditioning was performed as described above. The next day, animals underwent the fear recall test drug free.

### Optogenetic experiments

For bilateral optogenetic manipulations during behavior, the implanted optical fibers (200 μm core diameter, 0.22 NA, Thorlabs) were connected to 200 μm patch cords (Thorlabs) with zirconia sleeves and the patch cords were connected to a light splitting rotary joint (FRJ 1x2i, Doric Lenses) that was connected to a laser with a 200 μm patch cord (Thorlabs). For mice expressing the light-activated excitatory opsin ChR2 (ChR2-EYFP) and their EYFP controls, blue light pulses were delivered from a 473 nm laser (LuxX473, Omicron). Laser power at the tip of the optic fiber was 5-10 mW. DA terminals in the TS were excited at the time of the CS (CS-Paired) or the US (US-Paired) and TS neuronal activity was excited at the time of the US (US-Paired) during fear conditioning. For CS-Paired animals, 5-ms light pulses were delivered at 20 Hz during the CS presentation. For US-Paired animals, 5-ms light pulses were delivered at 20 Hz for 3s. The laser was turned on 1s before the US onset to 1s after the US offset.

### Histology

At the end of the experiments, mice were deeply anesthetized with sodium pentobarbital and were transcardially perfused with 4% paraformaldehyde and 15% picric acid in phosphate-buffered saline (PBS). Brains were removed, post-fixed overnight and coronal brain slices (60 µm) were sectioned using a vibratome (VT1000S, Leica). Standard immunohistochemical procedures were performed on free-floating brain slices. Briefly, sections were rinsed with PBS and then incubated in a blocking solution (10% horse serum, 0.5% Triton X-100 and 0.2% BSA in PBS) for 1 h at room temperature. Slices were then incubated in a carrier solution (1% horse serum, 0.5% Triton X-100 and 0.2% BSA in PBS) containing the primary antibody overnight at room temperature. The next day, the sections were washed in PBS and then incubated in the same carrier solution containing the secondary antibody overnight at room temperature. The following primary antibodies were used: polyclonal rabbit anti-tyrosine hydroxylase (TH, catalog # 657012, 1:1000, Calbiochem), monoclonal mouse anti-TH (catalog # MAB318, 1:1000, Millipore), polyclonal guinea pig anti-TH (catalog # 213004, 1:1000, Synaptic Systems), polyclonal rabbit anti-GFP (catalog # A11122, 1:1000, Life Technology), polyclonal chicken anti-GFP (catalog # AB13970, Abcam), and mCherry monoclonal anti-rat (catalog # M11217, 1:1000) Invitrogen). The following secondary antibodies were used: Alexa Fluor 568 goat anti-rabbit (catalog # A11011, 1:1000, Thermo Fisher Scientific, Invitrogen), Alexa Fluor 568 goat anti-mouse (catalog # A11004, 1:1000, Thermo Fisher Scientific, Invitrogen), Alexa Fluor 568 goat anti-guinea pig (catalog # A11075, 1:1000, Invitrogen), Alexa Fluor 405 goat anti-rabbit (catalog # A31556, 1:750, Invitrogen), Alexa Fluor 488 goat anti-rabbit (catalog # A11008, 1:1000, Thermo Fisher Scientific, Invitrogen), Alexa Fluor 488 goat anti-chicken (catalog # AB150173, 1:1000, Abcam), Alexa Fluor 568 goat anti-rat (catalog # A11077, 1:1000, Invitrogen). For DAPI staining, sections were incubated for 10 min in 0.1 M PBS containing 0.02% DAPI (catalog # D1306, Molecular Probes, Invitrogen). Finally, all sections were washed with PBS, mounted on slides embedded with a mounting medium for fluorescence (VECTASHIELD®, Vector Laboratories) and coverslipped.

### Statistics

Data were statistically analyzed using GraphPad Prism (GraphPad Software) and MATLAB (Mathworks). All statistical tests were two-tailed and had an α level of 0.05. All error bars show s.e.m. All ANOVAs were followed by Bonferroni *post hoc* tests if significant main or interaction effects were detected. No statistical methods were used to predetermine sample size, but our sample sizes were similar to those generally used in the fear conditioning field. Animals were randomly assigned to experimental groups before the start of each experiment after ensuring that all experimental groups were matched for age. For 6-OHDA lesioning, chemogenetic and optogenetic experiments, experimental and control groups were matched from littermate mice. All results were obtained using groups of mice that were run in several cohorts.

## Acknowledgements

We would like to thank Jochen Roeper for his support; Beatrice Fischer, Jasmine Sonntag, Sebastian Betz, Günther Amrhein and Thomas Wulf for technical assistance; and Torfi Sigurdsson for helpful discussions. This work was supported by the Deutsche Forschungsgemeinschaft (DFG Grant DU 1433/5-1 to S.D.).

---

**Supplementary Figure 1.**
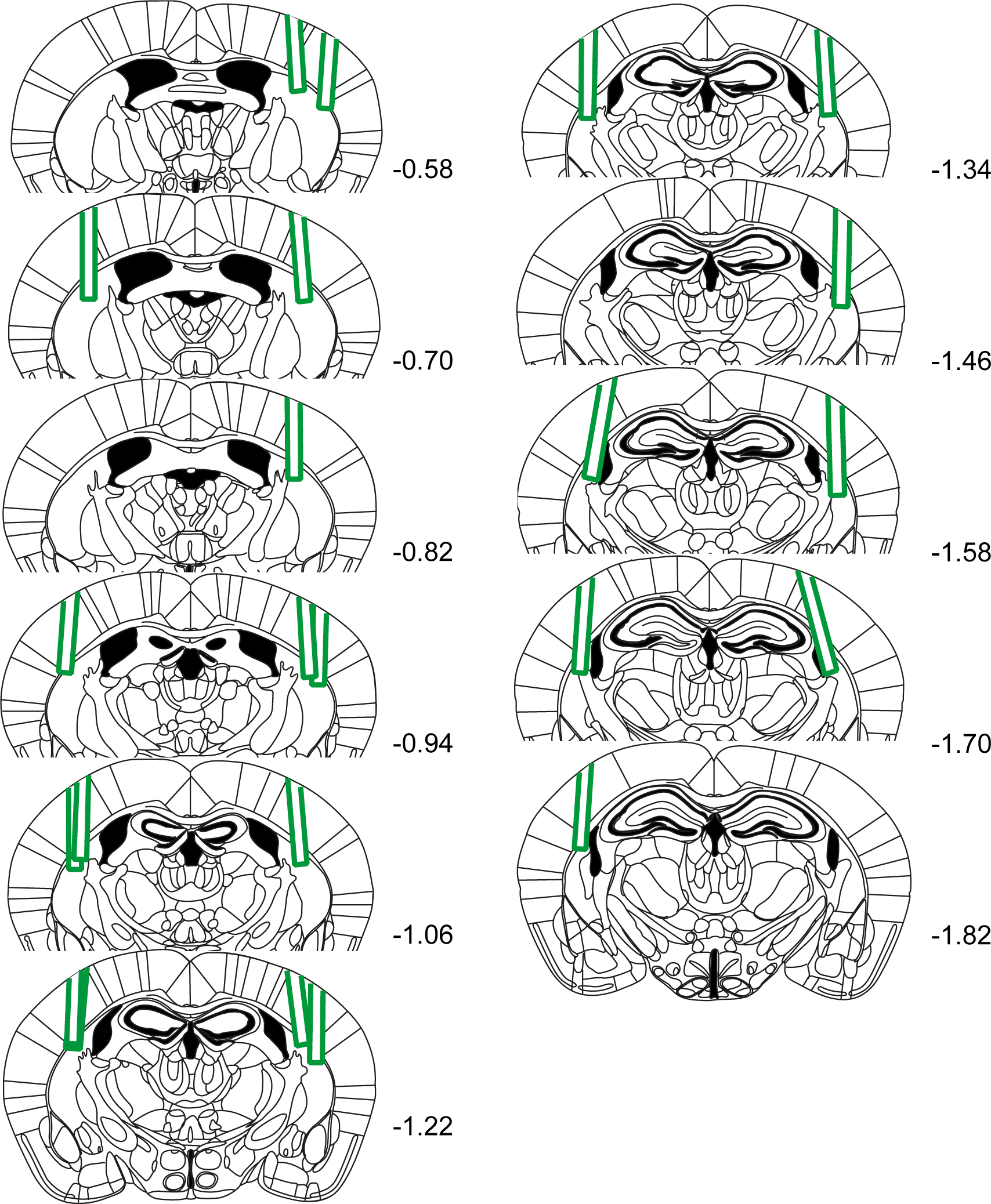
Optical fiber placements for GCaMP recordings of DA terminals in TS. Schematic coronal sections showing placement of optical fibers in TS. Numbers represent distance to bregma.

**Supplementary Figure 2.**
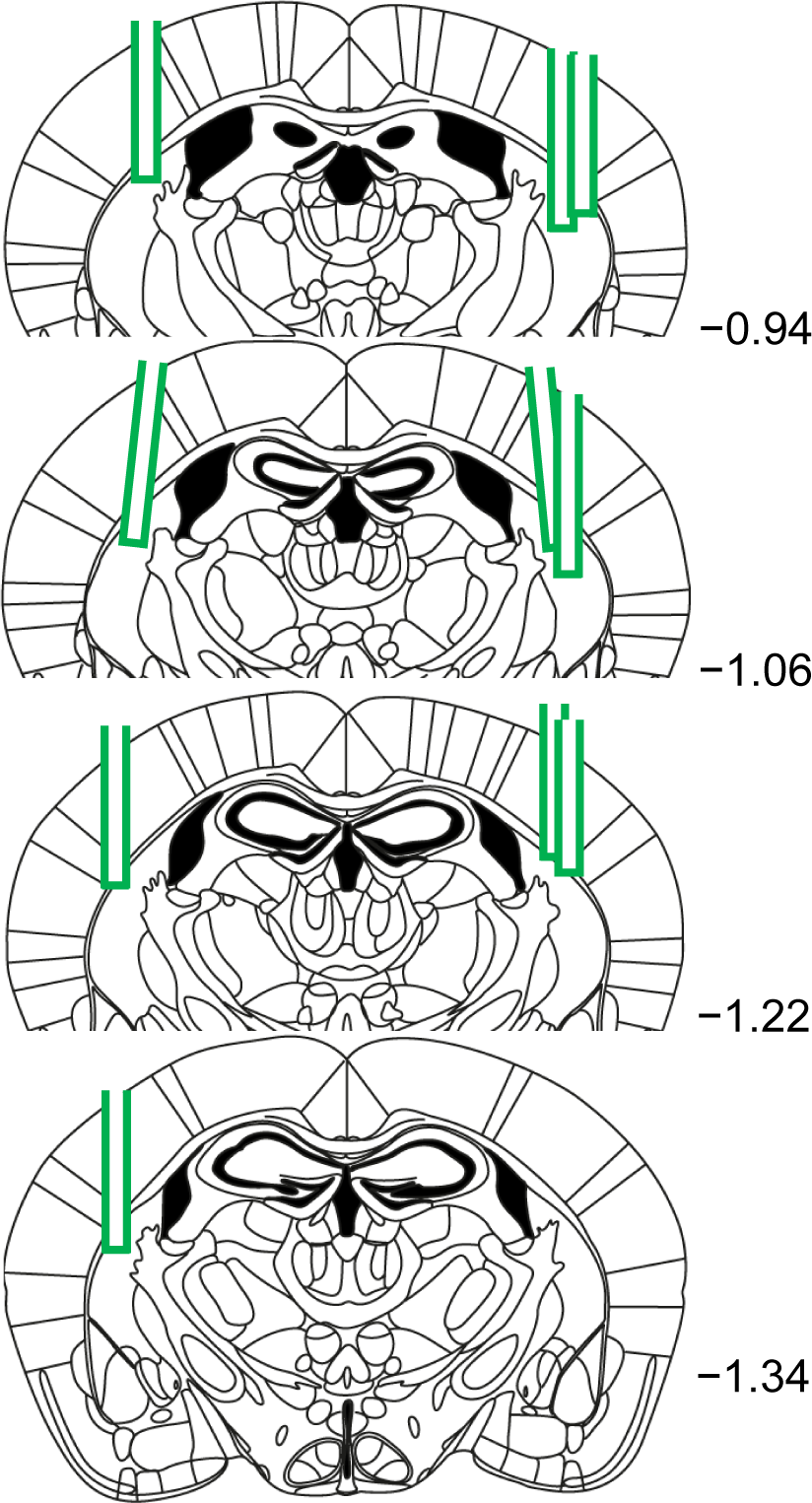
Optical fiber placements for dLight recordings in TS. Schematic coronal sections showing placement of optical fibers in TS. Numbers represent distance to the bregma.

**Supplementary Figure 3.**
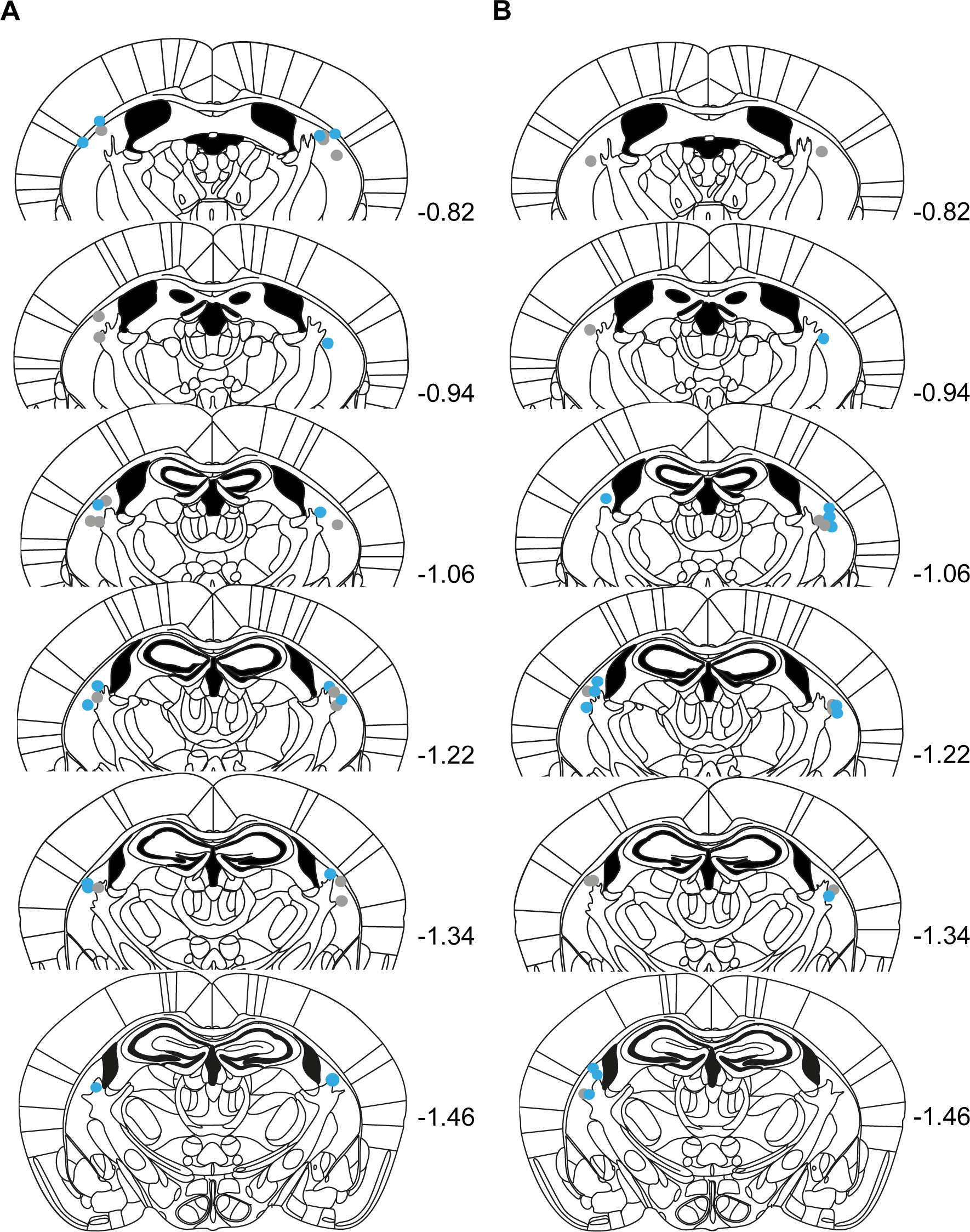
Optical fiber placements for optogenetic excitation of DA terminals in TS. Schematic coronal sections showing placement of optical fiber tips in TS for US-Paired ChR2 (blue) and US-Paired EYFP (gray) groups **(A)** and CS-Paired ChR2 (blue) and CS-Paired EYFP (gray) groups **(B)** . Numbers represent distance (mm) to bregma.

**Supplementary Figure 4.**
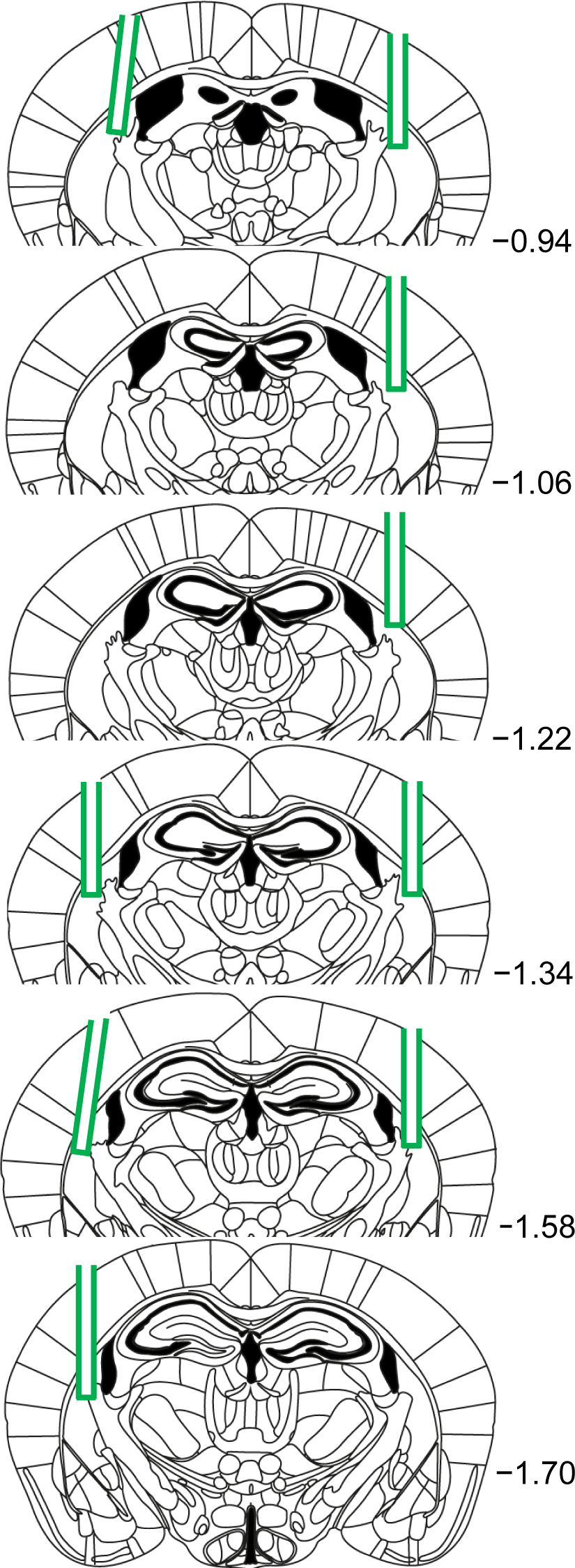
Optical fiber placements for GCaMP recordings in TS. Schematic coronal sections showing placement of optical fibers in TS. Numbers represent distance to the bregma.

**Supplementary Figure 5.**
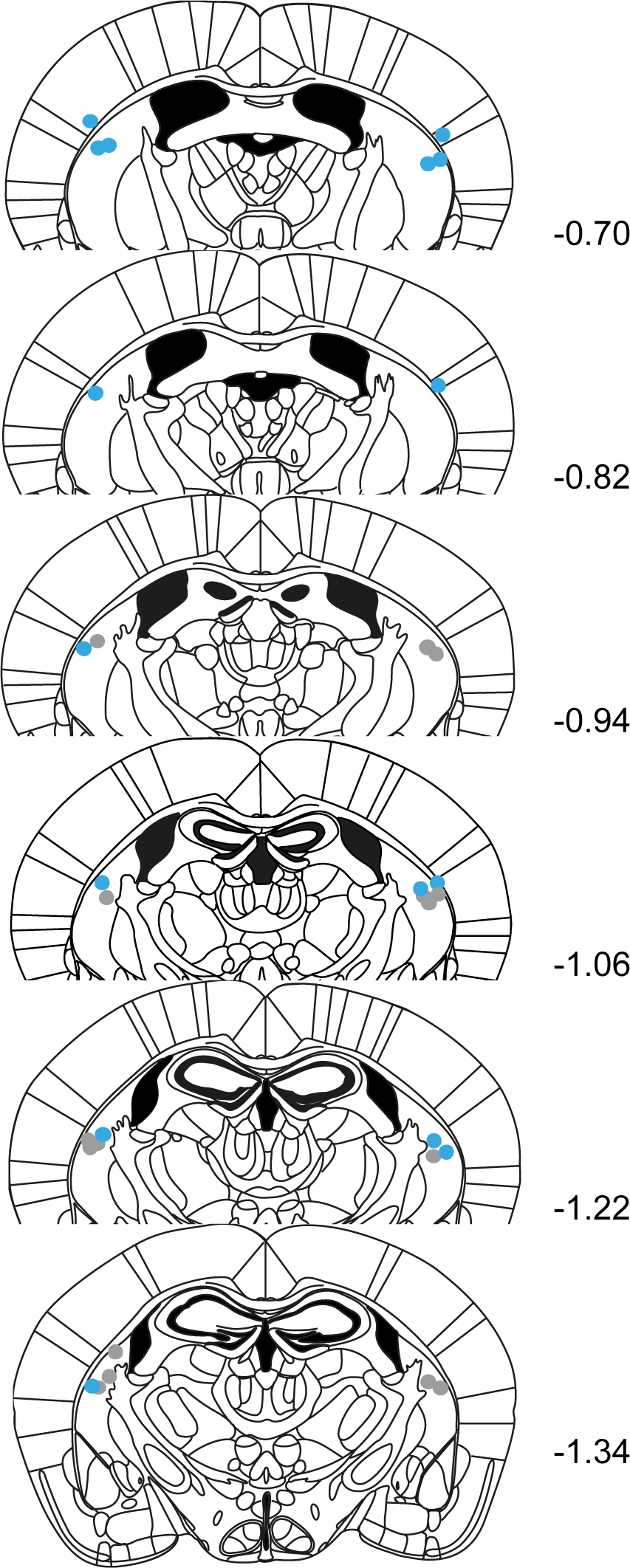
Optical fiber placements for optogenetic excitation of TS neurons. Schematic coronal sections showing placement of optical fiber tips in TS for ChR2 (blue) and control (gray) group. Numbers represent distance to bregma.

**Supplementary Figure 6.**
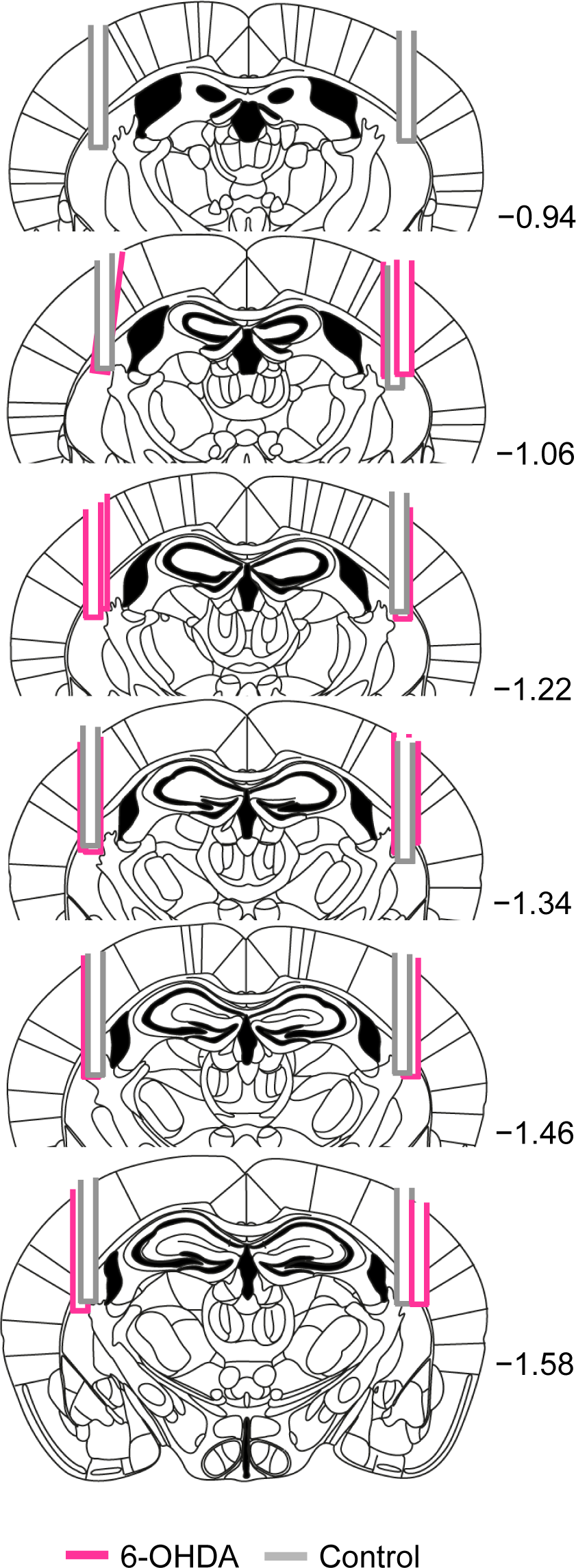
Optical fiber placements for GCaMP recordings in the TS following 6-OHDA Lesions of TS-projecting DA neurons. Schematic coronal sections showing placement of optical fibers in the TS for 6-OHDA (pink) and Control (gray) groups. Numbers represent distance to the bregma.

